# Recent origin of an XX/XY sex-determination system in the ancient plant lineage *Ginkgo biloba*

**DOI:** 10.1101/517946

**Authors:** He Zhang, Rui Zhang, Xianwei Yang, Kai-Jie Gu, Wenbin Chen, Yue Chang, Qiwu Xu, Qun Liu, Yating Qin, Xiaoning Hong, Yin, Inge Seim, Han-Yang Lin, Wen-Hao Li, Jinfu Tian, Shanshan Li, Liu, Junnian Liu, Shanshan Liu, Xiaoshan Su, Congyan Wang, Fu-Ming Zhang, Song Ge, Cheng-Xin Fu, Simon Ming-Yuen Lee, Yiji Xia, Jian Wang, Huanming Yang, Guangyi Fan, Xun Xu, Yun-Peng Zhao, Xin

## Abstract

Sexual dimorphism like dioecy (separate male and female individuals) have evolved in diverse multicellular eukaryotes while the molecular mechanisms underlying the development of such a key biological trait remains elusive (*1*). The living fossil *Ginkgo biloba* represents an early diverged lineage of land plants with dioecy. However, its sex-determination system and molecular basis have long been controversial or unknown. In the present research, we assembled the first and largest to date chromosome-level genome of a non-model tree species using Hi-C data. With this reference genome, we addressed both questions using genome resequencing data gathered from 97 male and 265 female trees of ginkgo, as well as transcriptome data from three developmental stages for both sexes. Our results support vertebrate-like XY chromosomes for ginkgo and five potential sex-determination genes, which may originate ~14 million years ago. This is the earliest diverged sex determination region in all reported plants as yet. The present research resolved a long-term controversy, lay a foundation for future studies on the origin and evolution of plant sexes, and provide genetic markers for sex identification of ginkgo which will be valuable for both nurseries and field ecology of ginkgo.

## MAIN TEXT

Dioecy (separate male and female individuals) and sex chromosomes are common in animals, but rare in plants (*2*). Plants appear to have sex-determination systems developed at different evolutionary timeslines, whereas the best studied animal systems are ancient. The repeated independent evolution of sex chromosomes in plants provide a unique system studying the time course of evolutionary events in sex chromosomes (*3*). Sex chromosomes in most flowering plants (angiosperm) are cytologically homomorphic or have very small non-recombining sex-determination regions on homomorphic sex chromosomes (*4–6*). Remarkable progresses have been achieved on understanding sex-determination in plants by applying genomic approaches in papaya (*Carica papaya*), cucumber (*Cucumis sativus*), and white campion (*Silene latifolia*) (*2*). In contrast, sex chromosomes are heteromorphic in all the six known species (0.6%) from three families (Cycadaceae, Ginkgoaceae and Podocarpaceae) in gymnosperm (plants with naked seeds), possibly reflecting an ancient evolutionary history (*3*). Unfornately, the confirmation of sex chromosomes and reconstruction of their evolutionary history remain to be explored in the sister lineage of flowering plants.

The dioecious maidenhair tree, *Ginkgo biloba* (ginkgo), a long-lived tree species which represents one of the four gymnosperm lineages, provides an ideal model for the origin of dioecy in seed plants. Gingko is an ancient lineage present in the Jurassic period 170 million years ago (*7*). The discovery of multiflagellated swimming sperm from a male ginkgo tree in 1896 hinted at a striking example of convergent evolution (*8*). Its sex-dermination system, XX/XY (*9, 10*) vs. ZW/ZZ (*11, 12*), remains controversial and was based on previous work using optical microscope or Fluorescence *in Situ* Hybridization (FISH) approaches, which may suffer unreliability or ambiguity (*9-11, 13*). Thus, the genetic basis of the sex determination of ginkgo has remained poorly explored (*2, 9, 11, 12, 14*) largely due to its large and complex genome (*15*).

We used three-dimensional proximity information obtained by chromosome conformation capture sequencing (Hi-C) to order and orient the draft genome assembly of *Ginkgo biloba* (*15*) (**fig. S1**), generating 12 chromosomes spanning 9.03 Gb (~94% of the whole genome) **(tables S1, S2**). The chromosome-level assembly was well supported by the inter- and intra-chromosome contact matrices of the Hi-C data, and revealed an anti-diagonal pattern within a chromosome and clear separations between chromosomes (**fig. S2a**). Moreover, chromosome sequence lengths were highly correlated with the physical length of chromosomes described in a previous karyotype study (*13*) (*R*^2^=0.98, **fig. S2b**), further supporting the high quality of our assembly.

We performed a genome-wide association study (GWAS) of ~1.5 million SNPs in 97 male and 265 female specimens sequenced at an average depth of ~8× (**table S3**; described in our parallel study (*16*): Zhao *et al.*, submitted simultaneously). A ~4.6 Mb region on chromosome 2 (Chr2: 380.00 Mb–384.60 Mb) was identified as a candidate sex-determination region (SDR) due to its significant association with sex (**Fig. 1A** and **fig. S3**, corrected *P*-value < 0.05). We computed the fixation index (*F*_ST_) (*17*) between the males and females in the whole genome, showing substantially higher differentiation on the similar region of chromosome 2 (**Fig. 1B**). Finally, the linkage disequilibrium (LD) of this region was substantially stronger than for other regions (**Fig. 1C** and **fig. S4**), consistent with the previous reports of enhanced LD on sex chromosomes in other species(*18*). To further validate this region as an SDR, six randomly-selected SNPs were sequenced in 24 males and 24 females using PCR and Sanger sequencing, revealing a high correlation between the SNPs and sexes (**fig. S5**). One SNP for PCR validation matched amplification products in 70.83% males with no product in all females (**Fig. 1D**). Using 19,164 SDR-associated SNPs, 97 males and 265 females were completely clustered into two distinct groups, and the remaining 183 sequenced ginkgo individuals with unknown sex information were classified into the two sex clusters, i.e., 75 males and 108 females (**Fig. 1E**).

**Fig. 1.**
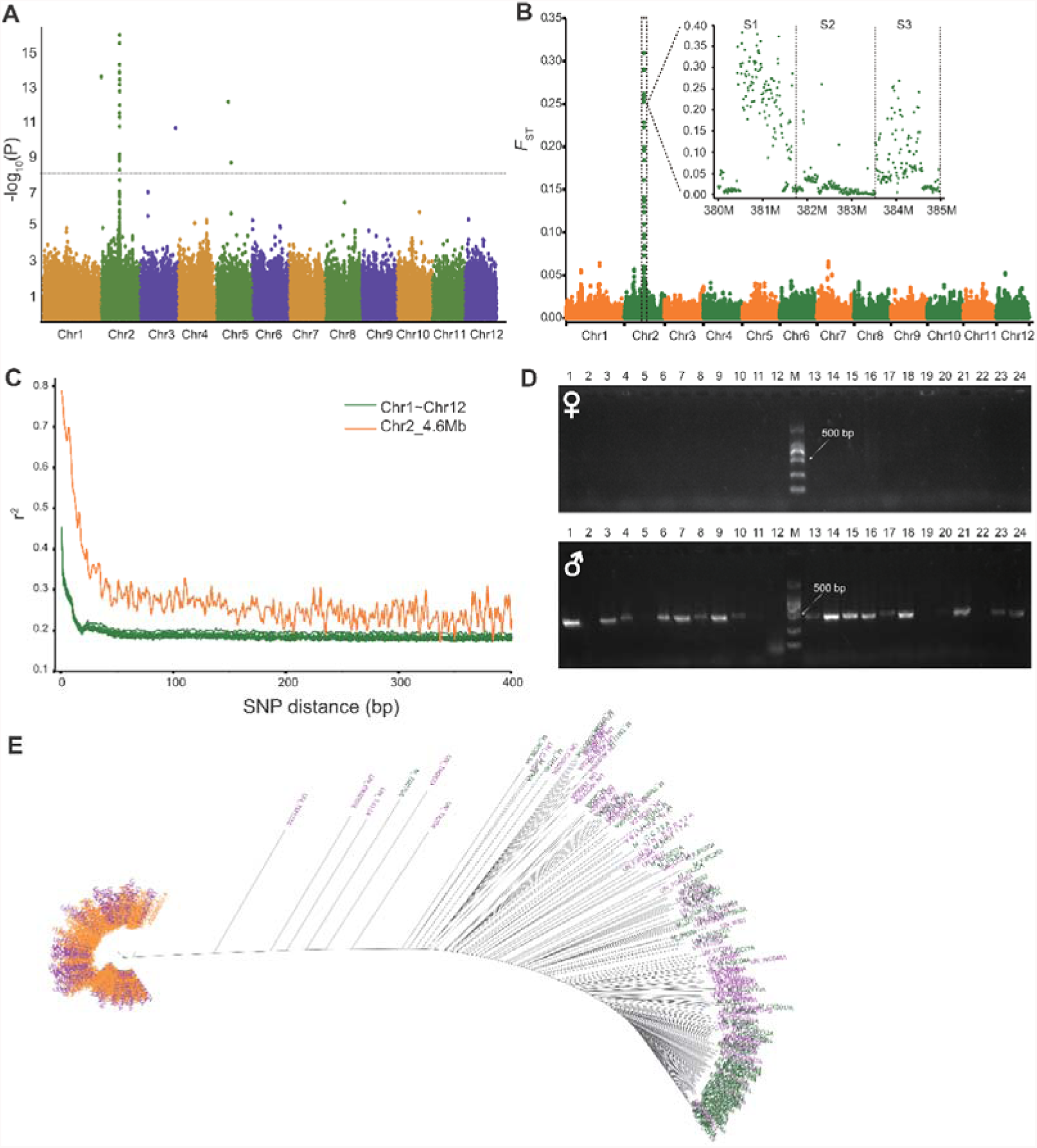
4.6 Mb region on chromosome 2 is a candidate ginkgo sex-determination region. **(A)** Manhattan plot of a genome-wide association study (GWAS) for SNPs associated with sex on 97 male and 295 female specimens. Negative log_10_ *P*-value from linear mixed model (*y*-axis) are plotted against SNP positions (*x*-axis) on each of the 12 ginkgo chromosomes. The genome-wide significant *P-value* threshold (10^-8^) is indicated by a horizontal line. **(B)** The 4.6 Mb region on Chr2 is highly differentiated between ginkgo sexes, indicated by high population differentiation (*F_ST_*) values (*x*-axis). Ginkgo chromosomes are shown on the *y*-axis. **(C)** SNPs spanning the 4.6 Mb region on Chr2 constitute a linkage disequilibrium (LD) block. Average genotypic association coefficient *r*^2^ (*y*-axis) is presented as a function of inter-SNP distance (*x*-axis) between 4.6 Mb region and whole genome. **(D)** Gel photograph of PCR amplicons. Validation of PCR amplification using specific designed primes for SNP on Chr2: 383,550,476 of 24 males and 24 females. **(E)** A neighbor-joining (NJ) tree constructed using 19,164 SNPs in the sex-determination region clusters 545 ginkgo specimen by sex. Specimens with known sex classification are labeled in orange (female; *n*=265) and green (males, *n*=97); specimens with unknown information (*n*=183) in purple.

Four male and five female accessions of ginkgo were resequenced at higher sequencing depth (~30×). Mapping the reads back to the reference genome continued to support the SDR (**figs. S6 and S7**). Males and females showed similar mapping depth (~32×) at both ends of the SDR (S1: 380.00–381.46 Mb and S3: 383.52–384.60 Mb) (**Fig. 2A**). The mapping depth in the middle of SDR (S2: 381.46–383.52 Mb) for males was ~17× (**Fig. 2A**), which was about half of that of females in S2 and also of males at the whole-genome level, indicating a higher divergence between males in S2. We observed notably higher SNP polymorphism in the S1 and S3 regions in males (97 males and 265 females) (**Fig. 2B**), and substantially higher heterozygosity (*P* = 5.84×10^-66^) in the S1 and S3 regions in the specimens subjected to deeper resequencing (~30×, four males and five females) (**Fig. 2C**). A ubiquitous feature of species with heteromorphic sex chromosomes (e.g. males with an X and a Y chromosome, females with two X chromosomes, as observed in mammals and fruit flies), is the presence of only one X chromosome in the male(*1*). Contrasting male and female, the copy number of S2 was one in males and two in females (**Fig. 2D**). The copy number of flanking regions (including S1 and S3) were two in both sexes(*19*). Collectively, we provide multiple lines of evidence of a classic XX/XY sex-determination system in ginkgo in which males harbor one allele (X) identical to the female alleles and one differentiated allele (Y; manifested by substantial divergence in SDR S2).

**Fig. 2.**
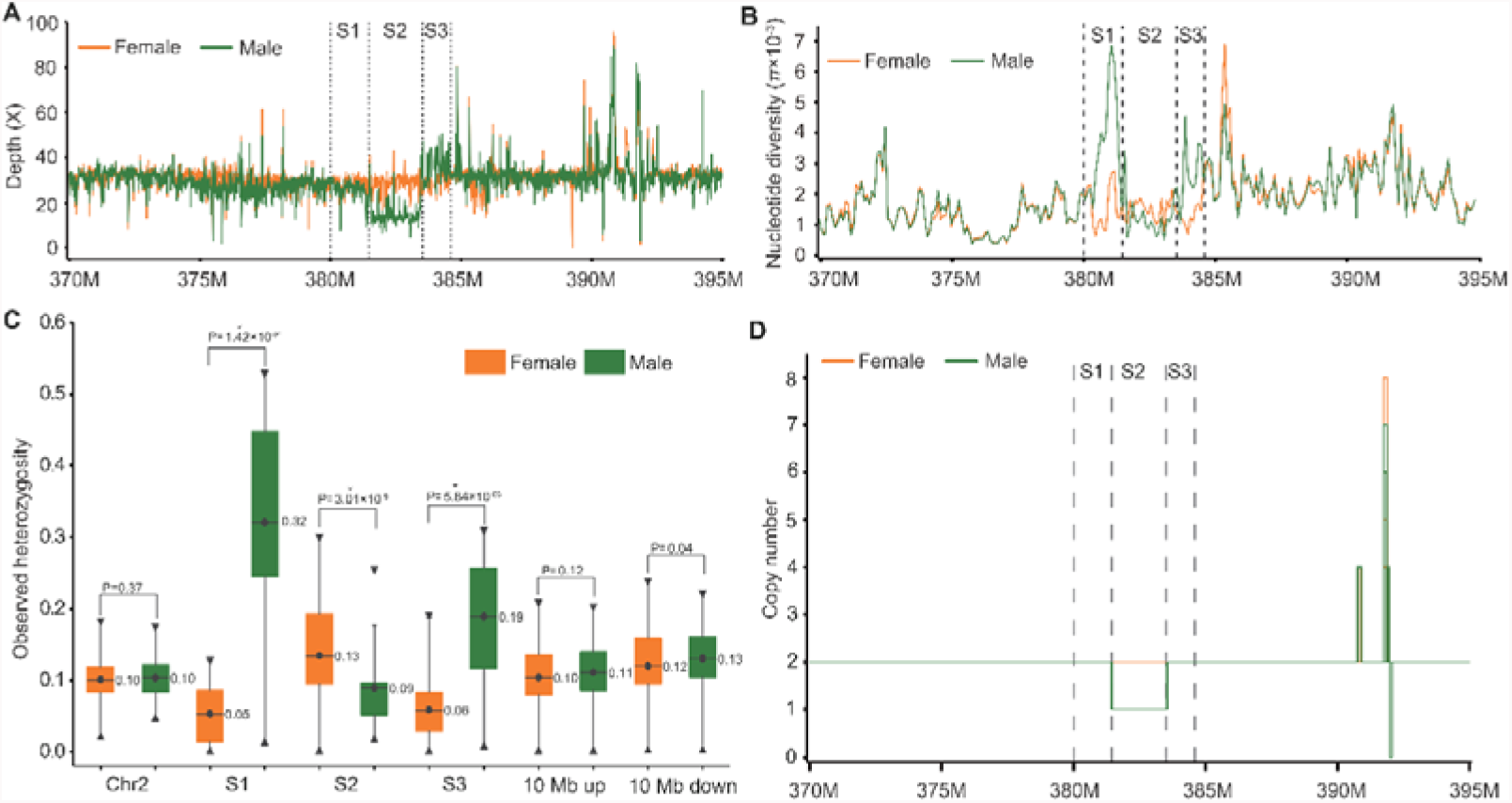
Further dissection of the ginkgo sex-determination region. Four males and five females were sequenced at high depth (~30×). **(A)** Sequencing depth in the sex-determination region is similar in both ends (sex-determination region 1, S1, Chr2:380.00-381.46 Mb; sex-determination region 3, S3, Chr2:383.52-384.60 Mb) but lower in males in the middle (sex-determination region, S2, Chr2:381.46-383.52 Mb). Average mapping depth of males and females are shown in 10-kb window. Depth on the *y*-axis, chromosome 2 region on the *x*-axis. **(B)** Nucleotide diversity (π) in the males are in the sex-determination region (especially in S1 and S3 regions). Examined using a 100-kb window size. Nucleotide diversity (π) on the *y*-axis, chromosome 2 region on the *x*-axis. **(C)** Observed heterozygosity in sex-determination regions S1 to S3 and chromosomal other regions in gingko sexes. Heterozygosity on the *y*-axis, chromosome region on the *x*-axis. **(D)** Estimated copy number is two for the S1 and S3 regions and one for the S2 region. Copy number on the *y*-axis, chromosome 2 region on the *x*-axis.

The origin of dioecy and sex chromosomes in ginkgo was inferred based on the three SDRs (S1, S2 and S3). We anticipated the highest difference between X and Y alleles in S2 because only the X allele could be assembled and reads from the Y allele could not be mapped back to this region. For S1 and S3, we had genotype data of both X and Y alleles, thus we calculated the pair-wise distances between four males and five females and estimated the divergence times of S1 and S3 to be 14.18 (11.04–16.64) million years ago (MYA) and 9.44 (4.48–11.42) MYA, respectively (**table S4**).

Considering the higher difference among X and Y alleles in S2, the divergence time of the associated SDR in gingko could be more ancient than 14.18 MYA. We were not able to reconstruct the most divergent part of the Y allele, possibly because of the inherent high complexity of the ginkgo genome. The emergence timing of ginkgo SDR, ~14 MYA, substantially precedes that for the known flowering plants, e.g., ~1.5 MYA for *Rumex hastatulus* with an XY sex-determination system, 5–10 MYA for *Silene latifolia*, and ~7.3 MYA for *Carica papaya*(*20*). Despite the oldest origin of ginkgo sex chromosomes in the known land plants to date, their timing as recent as ~14 MYA is in contrast to the ancient origin of ginkgo lineage and reproductive organs. Ginkgoalean and *Ginkgo* genus may have originated 320 MYA (*21, 22*) and 170 MYA (*7*), respectively. The timing for the origin of the extant species, *Ginkgo biloba*, was estimated 56–58 MYA (*23*). Fossils of ginkgo-like pollen and ovulate organs have long been known from the Upper Triassic and the Palaeozoic, respectively (*22*). Such a contrast again suggests a young origin of sex chromosomes in seed plants despite ancient origin of dioecy.

A critical property of an SDR is the presence of genes, typically transcription factors, associated with sexually dimorphic gene expression and traits (*1*). There were 16 protein-coding genes located in the ginkgo SDR (**Fig. 3A, Table 1** and **table S5**), among which five genes (*Gb_15883*, *Gb_15884*, *Gb_15885*, *Gb_15886*, and *Gb_28587*) were functionally annotated to be related to floral meristem development and sex determination. *Gb_15883* and *Gb_15884* were homologs of response regulators 12 and 2 (RR12 and RR2), proteins which respond to cytokinins and have been reported to be involved in the sex determination of kiwifruits (*24*). *Gb_15885* was a homolog of early flowering 6 (ELF6), a H3K4 demethylases gene which regulates flowering time (*25, 26*). *Gb_15886* was similar to Brassinosteroid-related AcylTransferase1 (AtBAT1) which regulates sex determination in maize (*27*). *Gb_28587* was a homolog of AGAMOUS-like 8, a member of a transcription factor family which specifies sex organ identity during development (*28*).

**Table 1.** Genes in the ginkgo sex-determination region (SDR). For each gene, the top-candidate ortholog, the associated SDR region, and allele specific gene expression is indicated (a dash indicates that no data could be obtained).

| Gene ID | Homologous<br>protein | Description | SDR | Allele |
| --- | --- | --- | --- | --- |
| *Gb_15883* | *RR12* | Response regulator 12 | S1 | X/Y |
| *Gb_15884* | *RR2* | Response regulator 2 | S1 | Y |
| *Gb_15885* | *ELF6* | ARLY FLOWERING 6 | S1 | Y |
| *Gb_15886* | *AT4G31910* | BR-related AcylTransferase1 | S1 | - |
| *Gb_28584* | *AT4G22030* | Putative F-box protein | S3 | X/Y |
| *Gb_28585* | - - |  | S2 | X/Y |
| *Gb_28586* | *SLO2* | SLOW GROWTH 2 | S2 | X/Y |
| *Gb_28587* | *AGL6* | AGAMOUS-like 6 | S2 | X |
| *Gb_28588* | *AT3G01930* | Major facilitator protein | S2 | X |
| *Gb_28589* | *AT4G31020* | Esterase/lipase<br>domain-containing protein | S2 | X |
| *Gb_30340* | - - |  | S3 | - |
| *Gb_30341* | *AT3G17450* | hAT dimerisation<br>domain-containing protein | S3 | - |
| *Gb_30342* | *AT3G17450* | hAT dimerisation<br>domain-containing protein | S3 | X |
| *Gb_30343* | *BIO1* | Biotin auxotroph 1 | S3 | - |
| *Gb_30344* | *AT4G30310* | FGGY family of carbohydrate<br>kinase | S3 | X |
| *Gb_30345* | *AT3G17450* | hAT dimerisation<br>domain-containing protein | S3 | - |

**Fig. 3.**
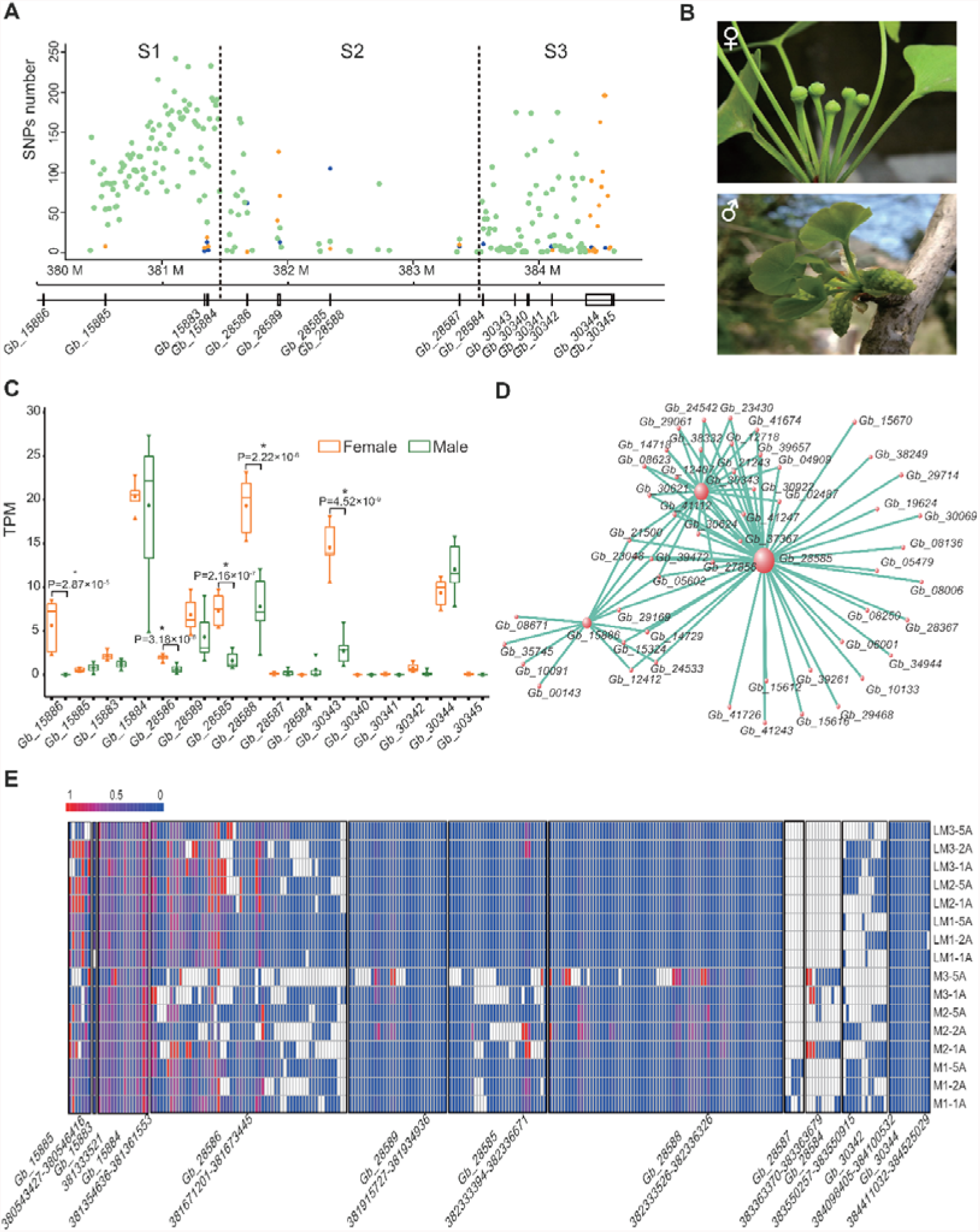
Identification of candidate ginkgo sex-determination genes. **(A)** Overview of 16 protein-coding genes in the sex-determination region. SNP counts on the *y*-axis, chromosome 2 region on the *x*-axis. **(B)** Images of male and female cones of ginkgo (photographed by Y.-P.Z.). **(C)** Differential expression of candidate sex-determination genes in ginkgo cone tissue. Expression (TPM, transcripts per million) on the *y*-axis, gene symbols on the *x*-axis. **(D)** Top 50 genes correlated with sex-determination genes in a ginkgo cone tissue co-expression network. Nodes represents genes and node size reflect the number of connections for each node. Three genes in the sex-determination region (Gb_15886, Gb_28585 and Gb_30343) were identified as hub genes (the most connected, or central genes). **(E)** A heat map showing the expression of 11 genes in sex-determination region with phased SNP haplotypes. 18 male samples are shown (8 cones, 8 leaves). Gene expression (TPM) shown on as a colour gradient: white being no expression detected; blue and red low and high Y haplotype expression, respectively.

To further determine the candidate sex-determination genes in ginkgo, we sequenced 32 transcriptomes from two tissues (cones and leaves, **Fig. 3B**) at various developmental stages in each of three males and females (**table S6**). Comparison between male and female specimens resulted in the identification of 5,831 and 132 differentially expressed genes (DEGs), respectively, in cones (reproductive organs) and leaves between male and female individuals (*P*-value < 0.001; **fig. S8a**, **b**). Of these 249 were uniquely expressed, specifically expressed genes (SEGs), in either male or female cones (**fig. S9** and **table S7**), while only two SEGs were found in leaves. Then, we focused on 5,774 genes which showed differential gene expression only in cones (**fig. S8c**). Five DEGs were located in the SDR, of which *Gb_15886* was exclusively expressed in female cones (**Fig. 3C**). To identify genes of S2 Y allele, we *de novo* assembled and contrasted the transcriptomes of males and females. This effort revealed five transcripts with significant divergence from the S2 X allele (~20% identity) which can be considered candidate Y allele-derived gene products. To understand the co-expression relationships between ginkgo genes, we performed weighted gene co-expression network analysis (WGCNA) (*29*) on male and female cone samples. This unsupervised and unbiased analysis identified a co-expression module significantly correlating 11 of 16 genes in the SDR with sex (**fig. S10** and **table S8**, *P*-value = 1×10^-6^). These genes (*Gb_15886*, *Gb_28585* and *Gb_30343*) were hubs, genes with the most connections with other genes in the network (**Fig. 3D**), further strengthening the identified SDR. Regulatory variants (SNPs) can also influence gene expression and provide a proxy of gene function (*30*).

We phased the X and Y haplotypes using the SNPs, resulting in 12,425, 2,067 and 4,672 SNPs phased in the S1, S2 and S3 with the average interval distances of 83 bp, 988 bp and 303 bp, respectively (**fig. S11** and **table S9**). Of these 19,164 phased SNPs, 17,533 were located in the intergenic regions and 1,361 in introns (**Fig. 3A**). Only 270 SNPs in 11 (out of 16) genes were located in the protein-coding exons. *Gb_15885* and *Gb_15884* specifically expressed Y allele genotypes, while the others nine genes expressed the X-allele or both X and Y alleles (**Fig. 3E** and **Table 1**). The allele-specific expression of *Gb_15885* and *Gb_15884* in male ginkgo might suggest that these genes have a critical role in sex determination analogous to *SRY* in mammals (*31, 32*).

In the present research, we identified the sex-determination region (SDR) of ginkgo and resolved its sex-determination system (XX/XY type). Our data revealed a gradually evolving and differentiating Y allele which originated approximately 14 MYA older than that in angiosperm. We provided evidence of a complex sex-determination process, including the expression of male and female sex-specific genes. Our genome and transcriptome data set offers an unprecedented resource for research on dioecy and sex chromosome evolution. In addition, we provided genetic markers for sex identification in vegetative stages of ginkgo (e.g., seedlings, juvenile trees, and non-flowering trees), which will be remarbly valuable for the selection of preferred sexes in nursery and for the identification of unknown sexes in field ecology.

## ACKNOWLEDGEMENTS

The authors are grateful to Dr. Qi Zhou for his constructive discussion and comments and to Fei-Da Ni, Lin-Jing Shi, Ya-Li Mu, Yi-Feng Lin and Ke Luo from Zhejiang University for their experimental assistance with validation of SDR. This study was supported by Shenzhen Municipal Government of China (JCYJ20151015162041454 to W. C. and JCYJ20150529150505656 to X. L), National Key R&D Program of China (No. 2017YFA0605104), the National Natural Science Foundation of China (No. 31870190) and the grant from the Chinese Academy of Sciences (CAS/SAFEA International Partnership Program for Creative Research Teams).

## Author contributions

X.L., G.-Y.F., Y.-P.Z. and X.X. conceived the project. Y.-P.Z., S.G., C.-X.F., S. M-Y. L., Y.-J.X., J.W., H.-M. Y. and X.X. supervised the study. Y.-P.Z., P.-P.Y., H.-Y.L. and W.-H.L. collected samples. K.-J.G conducted validation experiments. H.Z., R.Z., X.-W.Y., K.-J.G, W.-B.C., Y.C., Q.L., X.-N.H., J.-F.T., S.-S.L., L.-Q.L., F.-M.Z. and J.-N.L. conducted data analysis. C.-Y.W., X.-S.S. and S.-S.L. conducted the genome and transcriptome sequencing. Q.-W.X. and Y.-T.Q. constructed the Hi-C library and quality control. G.-Y.F., H.Z., X.L., Y.-P.Z., I.S. and R.Z. wrote the manuscript with the help from all co-authors.

## Data and materials availability

The ginkgo chromosome-level genome involved in this study was deposited in CNSA with the accession number CNA0000042 (Project ID: CNP0000122). All other relevant data supporting the findings of the study are available in this article, in the Extended Data, or from the corresponding authors upon request.

