## Supplementary Materials for "Recent origin of an XX/XY sex-determination system in the ancient plant lineage *Ginkgo biloba*"

**This PDF file includes:**

Materials and Methods
Figs. S1 to S11
Tables S1 to S9

### Materials and Methods

Hi-C library construction, sequencing and assembly

To get a high-resolution genome contact map, we used in situ Hi-C follow the protocol of previous study (*33*) with some modifications. The fresh plant leaves were crosslinked with 1% formaldehyde. To destroy the cell wall, formaldehyde fixed powder was added to Buffer solution. The restriction endonuclease MboI was used to digest DNA, followed by biotinylated residue labeling. The Hi-C library was then sequenced on BGISEQ-500 platform with 50 bp pair-end sequencing. HiC-Pro pipeline (*34*) was implemented in quality control. Of all 653,202,535 raw pair-end reads, there are 32% (207,324,555) paired Hi-C reads are valid and suitable for following analysis (**fig. S1**). Basing on these valid Hi-C reads, we used Juicer (*35*) and Aiden lab's Hi-C assembly pipeline (*36*) to assemble the genome with the main parameter "-m haploid -s 4 -c 12". As those tiny scaffold (shorter than 500bp) in original genome have fewer Hi-C contacts and difficult to analyze, we removed these scaffolds from our final assembly.

Genome-side association study (GWAS)

GAPIT (version 2) (*37*) was used to carry out the GWAS on sequencing data from 97 male and 265 female ginkgo specimens. Briefly, we extracted data of 97 males and 265 females from a VCF file of 545 individuals (Zhao *et al.*, under review). A total of 1,517,977 SNPs were retained. Three softwares and associated models were used to perform the GWAS analysis: GLM (general linear model) of PLINK (v1.90b4.5 64-bit), MLM (mixed linear model) of FastLmmC (v2.07.20140723), and cMLM (compressed mixed linear model) of GAPIT (v2017.08.18). The main findings from three software were unanimous, among which the results of GAPIT seemed to be better and is shown in **fig. S3**. The cMLM model can deal with bias caused by population structure and familial relatedness (kinship) and can be described as:

$$Y = X\beta + Zu + e,$$

where Y represents the phenotype; vector β is the fixed effects of population structure (Q); vector u explains the random effects of kinship (K) among individuals; vector e is unobserved factors; X and Z are the Henderson's matrices related to β and u.

Similar individuals were clustered into one group to reduce computational time.

To verify the credibility of the model, we use R package 'qqman' to draw the quantile-quantile plot **(**QQ-plot, **fig. S3a**). The QQ-plot showed that population structure and kinship effects were well controlled and provided evidence of the high reliability of GWAS results.

Depth, $F_{ST}$, SNP polymorphism and heterozygosity analysis

To determine if there is a significant difference in the sex-determination region (SDR) between males and females, we sequenced the genome of nine specimens (4 males and 5 females) at high depth. Briefly, reads were mapped against the reference genome using BWA (version 0.7.12-r1039) (*38*) and SAMtools (v1.8) (*39*), revealing a mean depth of ~30×. The sequencing depth distribution was calculated and plotted 10 kb windows. We calculated the genome-wide $F_{ST}$ between 97 males and 265 females with a 100 kb sliding window. The scatter diagram of $F_{ST}$ within and flanking a ~4.6 Mb candidate sex-determination region (SDR) in Chr2 was drawn. SNP polymorphism of females and the males was calculated using VCFtools (v1.15) (*40*) with a 100 kb sliding window. We also calculated observed heterozygosity of every individual in different regions using VCFtools (*40*). A Student's *t*-test was performed to detect whether there was a significant difference between males and females in S1, S2, S3 as well as other regions.

Haplotype phasing

A SNP can be phased if it satisfies the conditions that it is homozygous in all females and heterozygous in all males. In nine individuals with high sequencing depth (~30×; 4 males and 5 females), we obtained 19,164 phased SNPs from the 4.6 Mb candidate SDR. We next used these SNPs to calculate the 1-IBS matrix with PLINK (v1.9) (*41*), followed Neighbor-joining tree construction with MEGA (v7) (*42*) and tree figuration with FIGTREE (v1.43).

SNP phasing and Allele specific PCR

Primers were designed at positions upstream and downstream of the SNP at a distance of about 50 bp from the SNP to amplify SNP-rich fragments from 48 females and 48 males. The product sequence was sequenced using Sanger to confirm sex specificity of the SNPs in SDR. AS-PCR (allele-specific PCR) is to amplify specific bands for genotyping purposes. We designed specific primers for the SNPs in SDR. In addition, we added an extra mismatch at 4~5 bp from the 3' of the primer to improve specificity. We selected 24 males and 24 females to verify the efficacy of the probe. 20 pairs of primers were designed and tested. We filtered out 4 pairs of primers with a sensitivity greater than 0.65. Statistically, using 9 pairs of primers can reduce the theoretical probability of false negative results in male individuals to less than 1‱. All PCR experiments are divided into two parts: pre-experiment for figuring out the optimal temperature and validation experiments for verifying the efficiency. For pre-experiment,25μl of PCR mix (TsingKe® 1.1×T3 Super PCR Mix) is added into tubes each containing an individual's DNA and the following thermal cycle repeated 35 times after 3min of denaturation at 94℃:94℃—10secs, 48~60℃ (annealing temperature gradient) —10secs, 72℃—20secs. Except annealing temperature, the other reaction conditions of the validation experiments were the same. The amplification products are separated on a 3% TsingKe® agarose gel prepared in 1X TBE buffer and run in 1X TBE buffer.

Divergence time

All SNPs from the S1 and S3 regions were used to estimate divergence time between female and male ginkgo trees separately. Theoretically, the divergence time is equal to the value of twice the evolutionary rate divided by the genetic distance. According to the SNP data of 4.6 Mb candidate SDR and the reference genome, we extracted DNA sequences for the nine specimens with higher depth sequencing data (~30×; 4 males and 5 females) and employed these sequences as input to DISTMAT (EMBOSS:6.6.0.0) to calculate genetic distance.

RNA-seq /Differential expression/ Enrichment

Transcriptome of male and female cones (reproductive organs) and leaves were sequenced and used in differential gene expression analysis. We employed HISAT2 (v2.1.0) to index and map RNA-seq reads to the reference genome, SALMON (v0.9.1) to obtain normalized gene counts (transcripts per million; TPM), and STRINGTIE (v1.33) to detect differential gene expression in male and female tissues (*43*).

Allele-specific expression analysis

Based on the SNP calling, the distribution of SNPs in protein-coding, intron and inter-gene regions were visualized using a 10 kb window. We next mapped the transcriptome reads from 26 specimens to the ginkgo reference genome and counted the number of X- or Y- allele that supported each site using mpileup program in SAMtools(*39*) (v1.4). As in the female samples, nearly all the sites were only supported by Y-allele transcripts, we summarized the 16 out of 18 male samples (2 samples with poor mapping ratio were filtered). The dominant source of each site was then presented by the ratio of the X-allele evidences in a heat map.

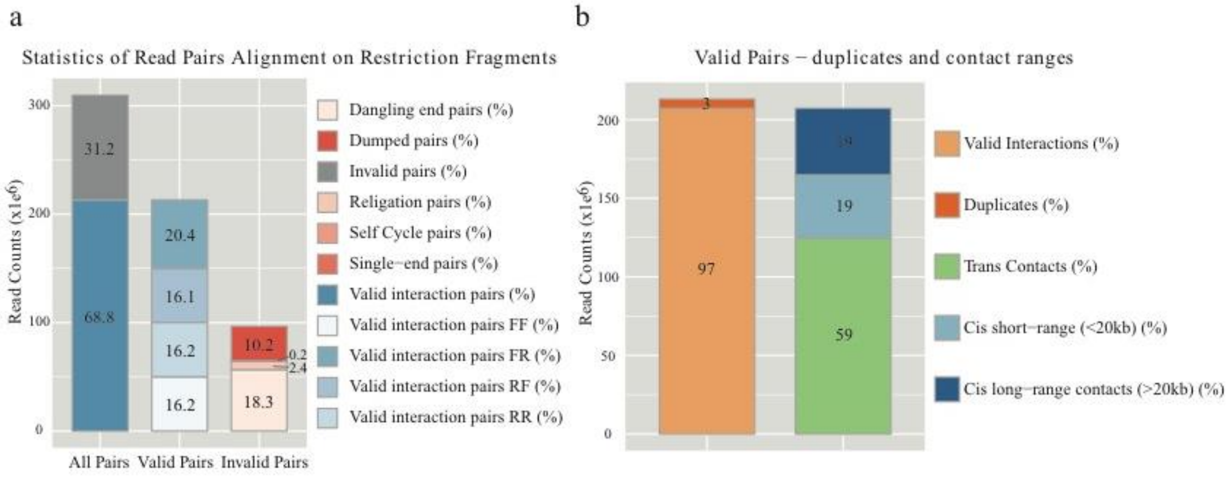


**Fig. S1. Quality control of chromosome conformation using Hi-C sequencing data of ginkgo (*Ginkgo biloba*). a.** Validation of the Hi-C reads according to the mapping results. For those 47.4% reads that can mapping to the reference genome (scaffolds), 68.8% reads were found to be valid (read pairs involve two different restriction fragments indicating chromosome conformation) and 31.2% were found to be invalid reads (reads pairs located on the same restriction fragment which resulted from duplications and etc.). The categories of different valid and invalid reads were also shown ('F' and 'R' referred to the direction of aligned reads, F for forward and R for reverse). **b.** The contact of the valid Hi-C reads. After removing the invalid, the valid pair-end reads were further categorized by contact range. Pair-end reads locate in the same and different scaffolds were *cis*- and *trans*-contacts, respectively. For the *cis*-contacts, short-range meant the distance of pair-end reads less than 20 kb.

a

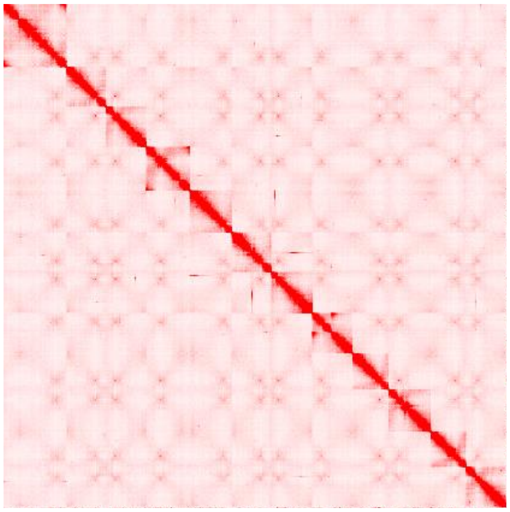

b

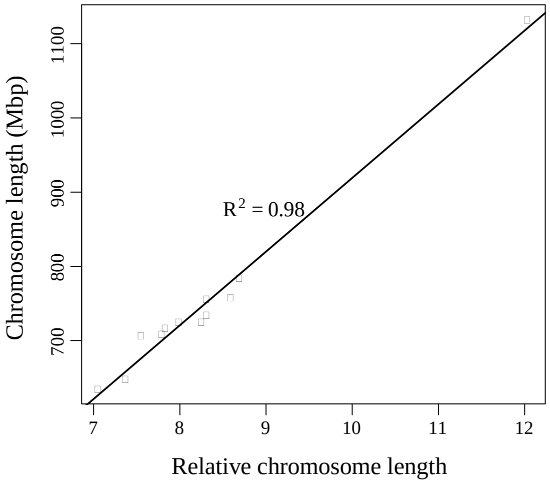


**Fig. S2. Chromosome-level assembly of ginkgo genome using Hi-C data. a.** Heatmap of contact metrics generate from mapping of Hi-C reads to genome sequences. **b**. Correlation coefficients between Hi-C assembled chromosome length and physical length.

a

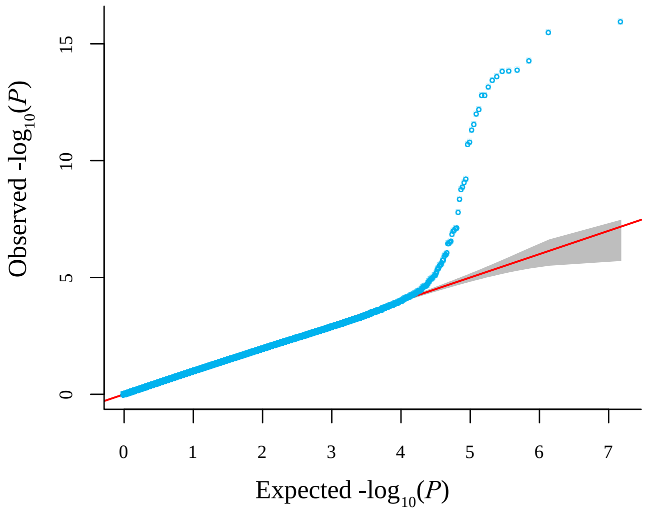


b

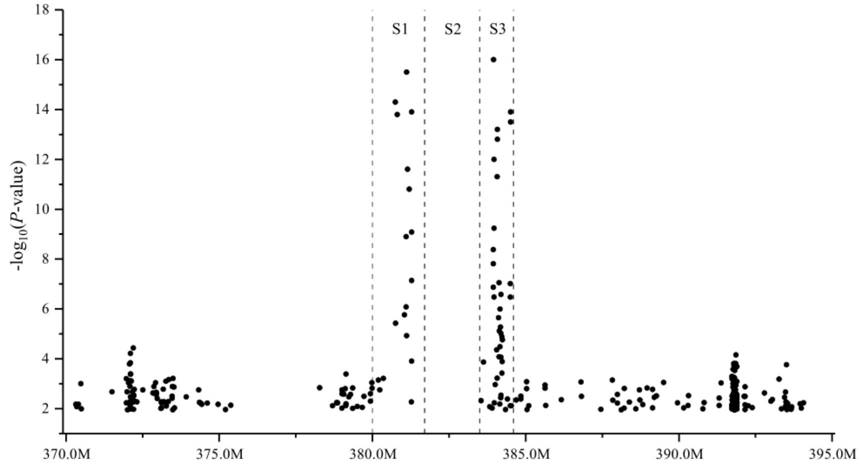


**Fig. S3. Genome-wide association study (GWAS) results for sex-related regions of ginkgo. a.** Q-Q plot for GWAS *P*-values of all single nucleotide polymorphisms (SNPs). **b.** Manhattan plot for GWAS *P*-values of all SNPs in a 25 Mb region of chromosome 2. The Manhattan graph shows that most of the SNPs associated with sex are located in a 4.6 Mb region on chromosome 2 (Chr2:380.00 Mb-384.60 Mb).

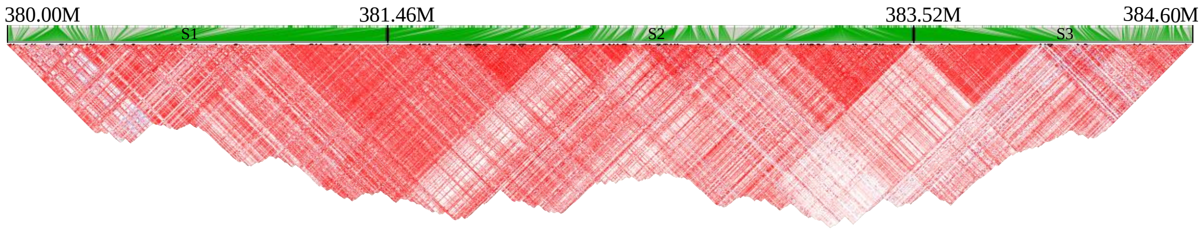


**Fig. S4. Linkage disequilibrium heatmap of females in SDR.** Only females were used for LD analysis, illustrating and proving that linkage disequilibrium in the SDR region is continuous.

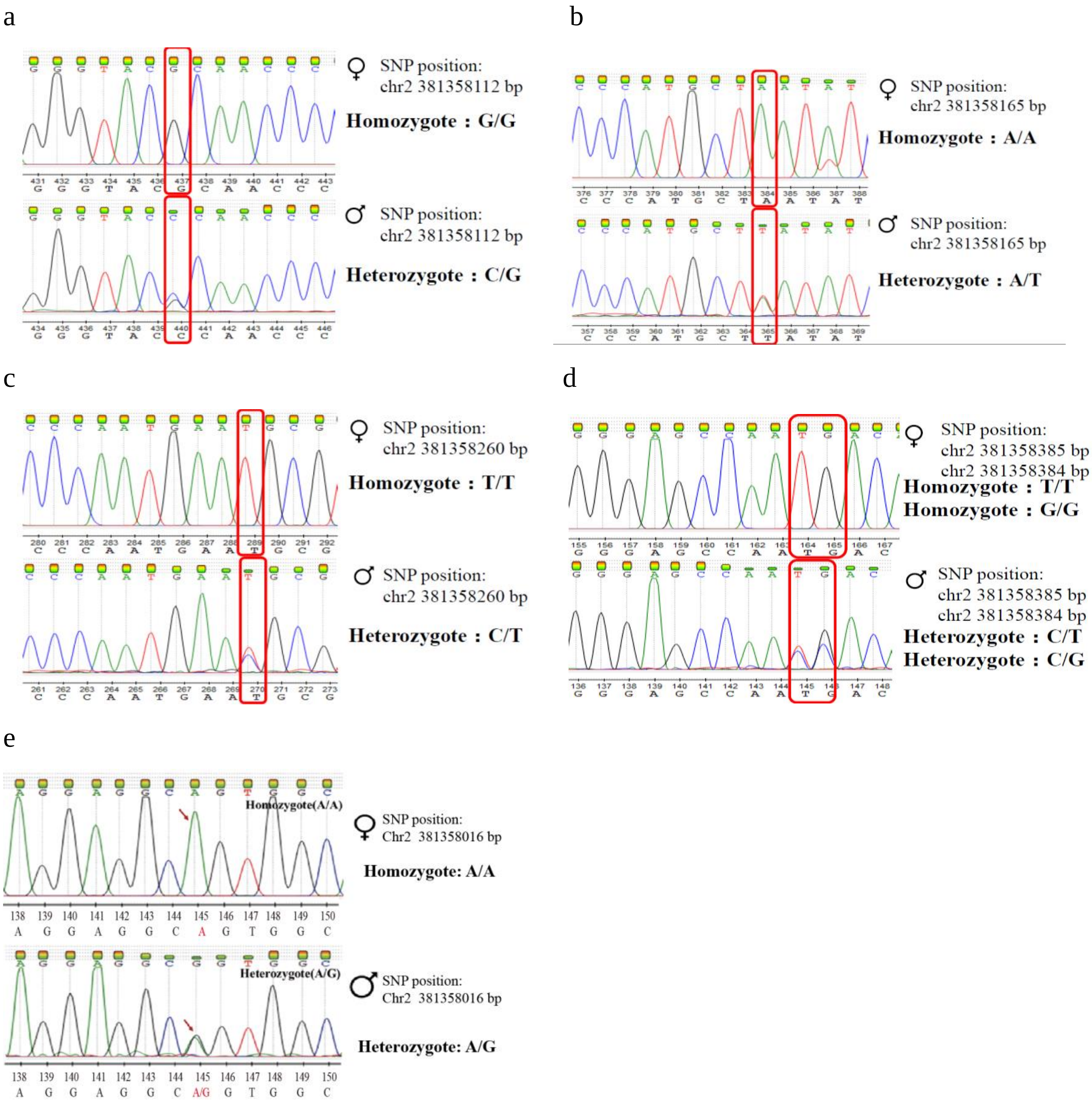


**Fig. S5. Validation of the sex-determination region (SDR) of ginkgo using five SNPs in 24 males and 24 females. a-e.** Overview of chromatograms of PCR products subject to Sanger sequencing. Single peak indicates that the genotype of the site is homozygous while double peaks means a heterozygous genotype.

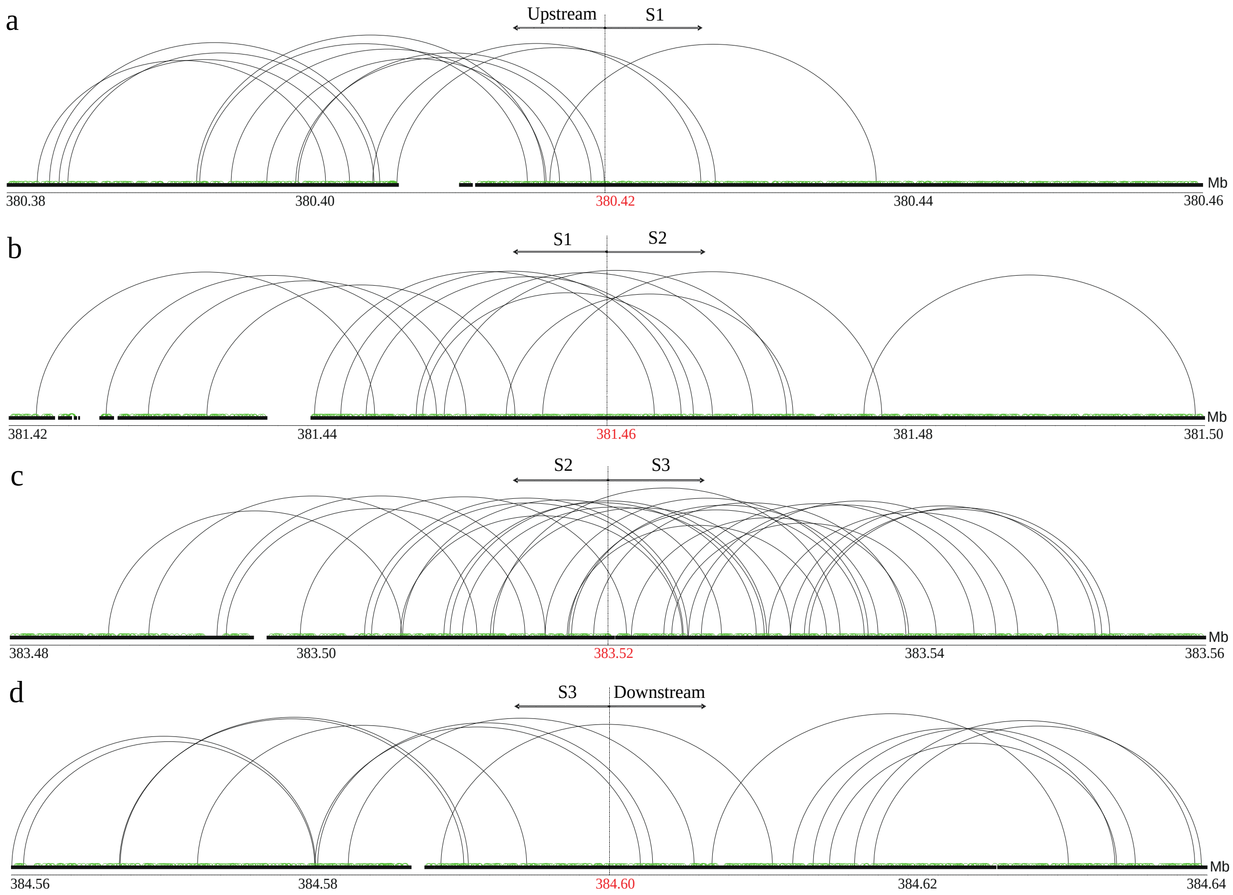


**Fig. S6. Pair-end reads mapping in sex-determination region (SDR) of ginkgo. a-d.** Mapping information of pair-end reads around each border of different sex-determination regions. Short insert-size library-derived reads (500bp) are colored in green; long insert-size library-derived reads (20 kb and 40 kb) in black. The coordinates of each border are colored in red.

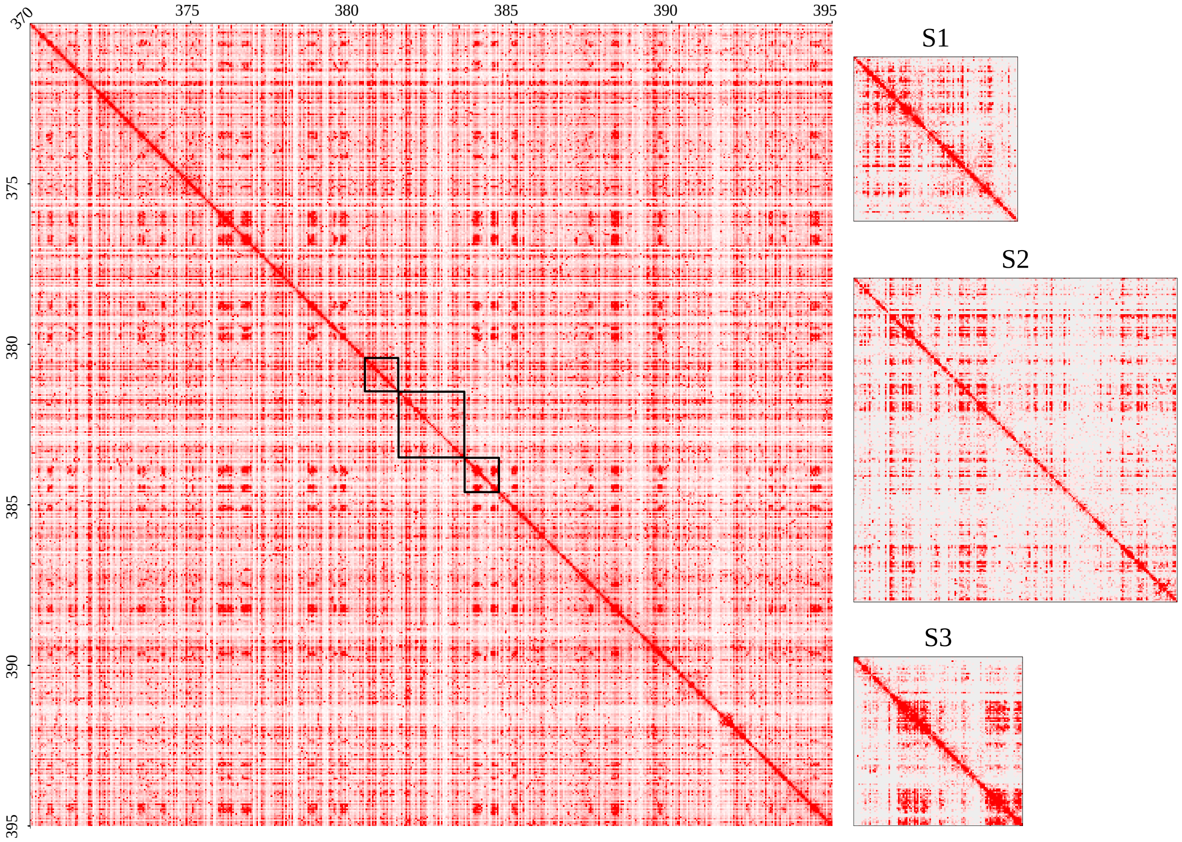


**Fig. S7. Partial contact matrix of ginkgo Hi-C data.** The left figure shows the contact matrix (contact information based on the reads) of chromosome 2, region 370-395 Mb, at 50 kb resolution. The three figures on the right show the sex-determination regions (SDR) S1 to S3 at a higher (10 kb) resolution.

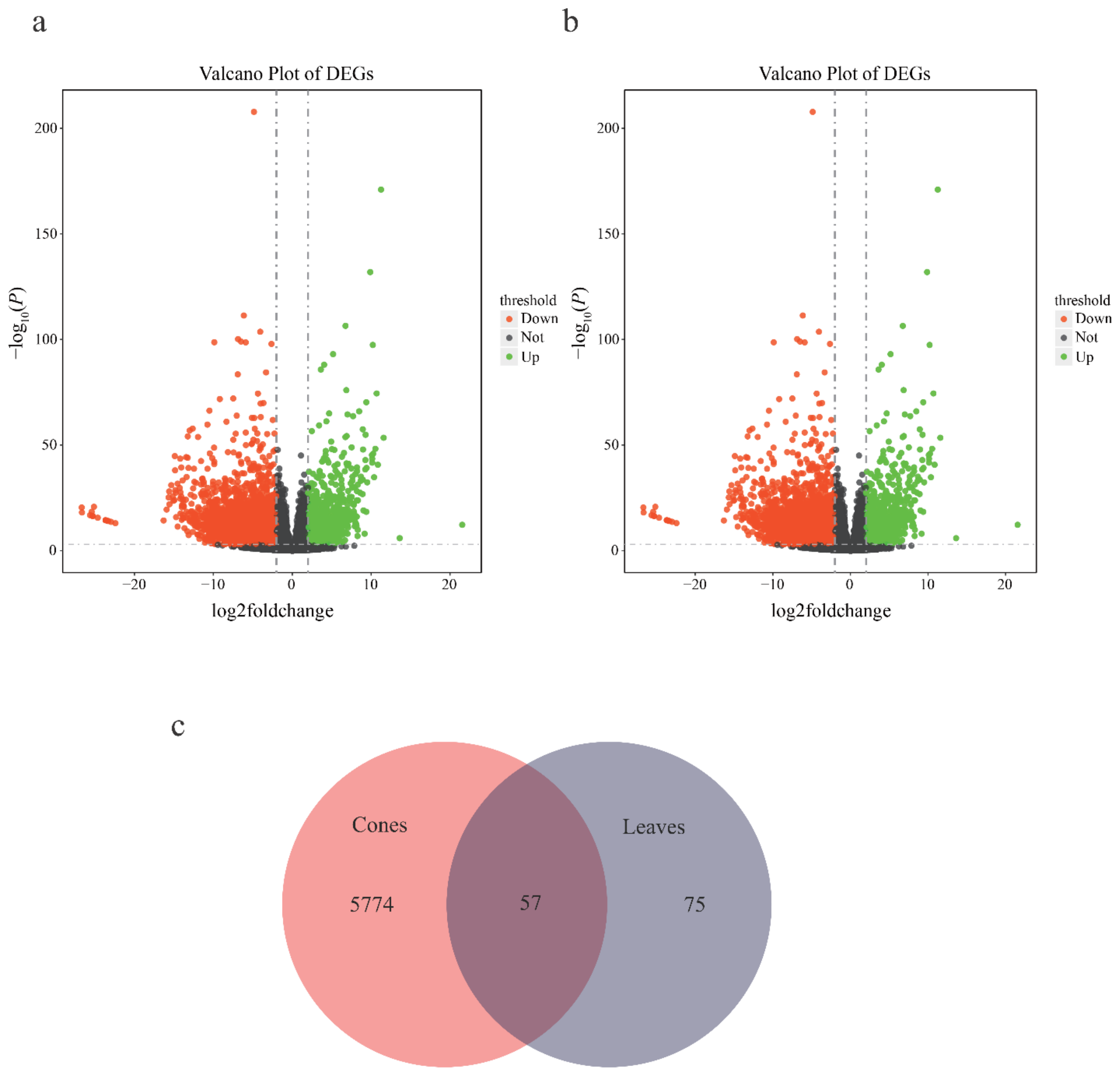


**Fig. S8. Differential expression genes (DGEs) between male and female ginkgo. a.** DEGs in cones (reproductive organs). Each dot indicates a gene; significantly up-regulated genes are shown in green, while significantly down-regulated genes are shown in red. **b.** DGEs in leaves. Annotated as in figure S8a. **c.** Venn diagram comparing DEGs in cones and leaves. 5,774 genes are DEGs in the reproductive tissue, cones.

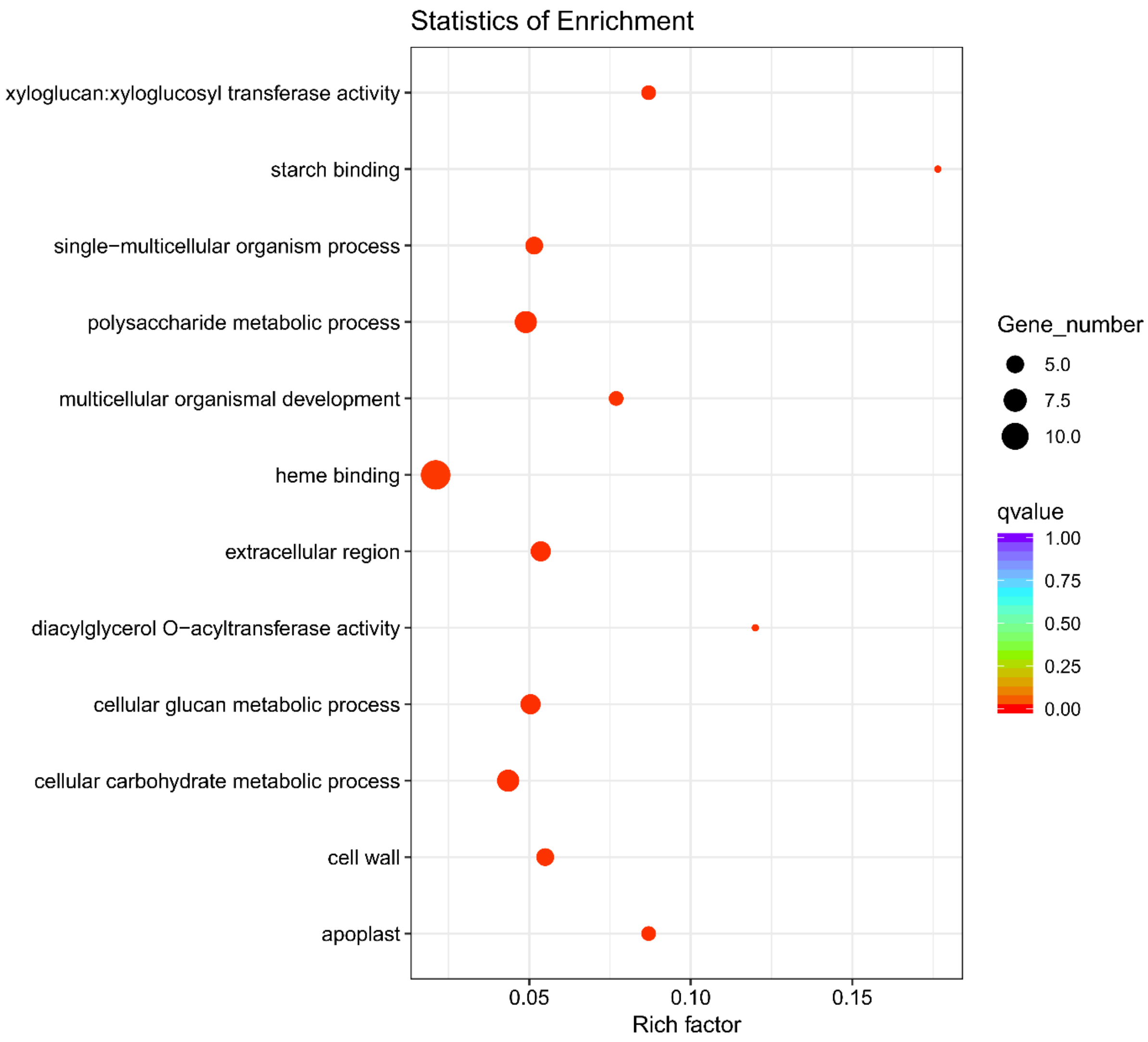


**Fig. S9. Functional enrichment of DEGs in male and female ginkgo cones.** DEGs between male cones and female cones are enriched in hormone signal transduction, flavonoid biosynthesis, and starch and sucrose metabolism.

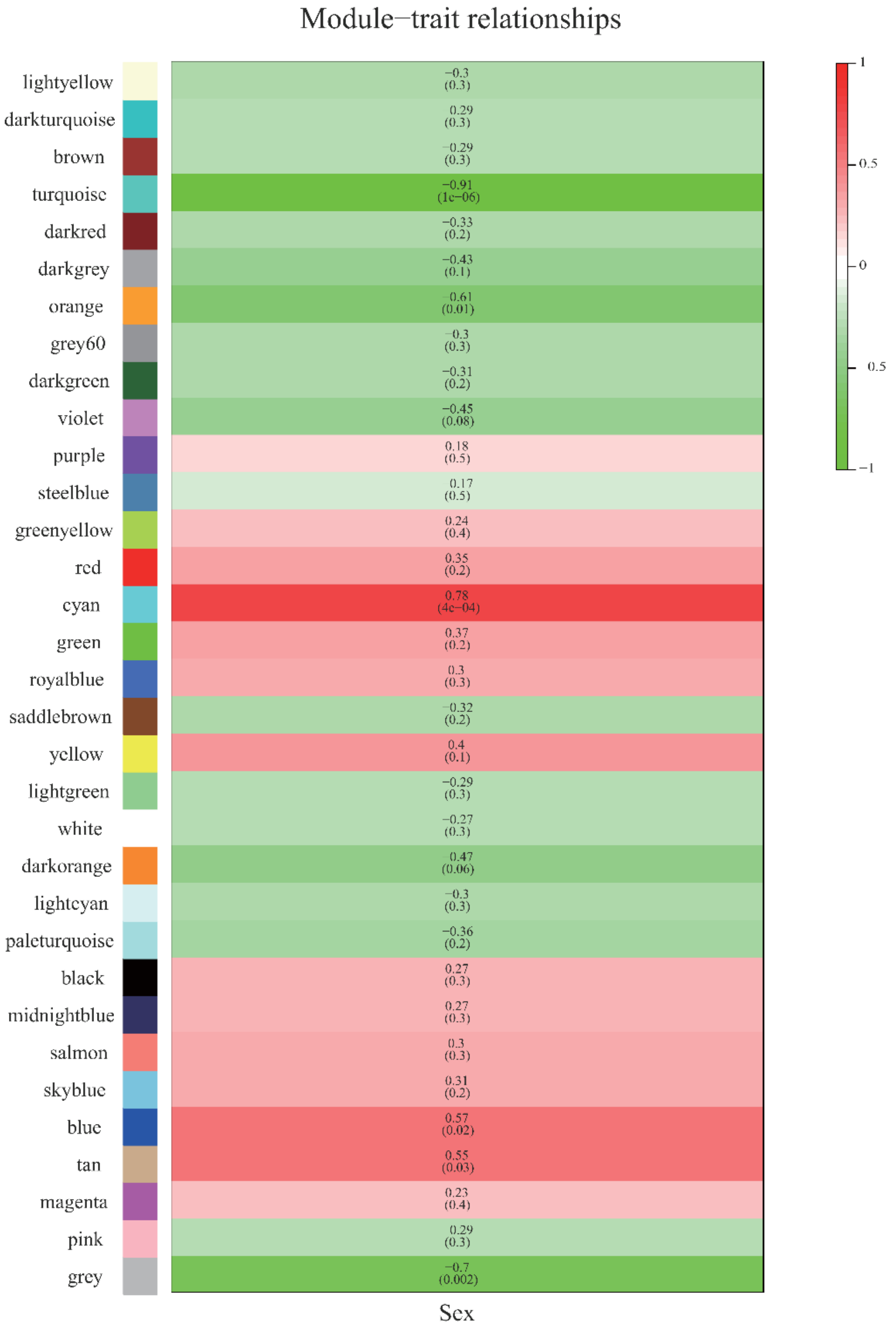


**Fig. S10. Network analysis of ginkgo cone transcriptomes identifies a co-expression module correlating with sex.** The bar on the left side represent distinct modules, each assigned a unique colour, identified in the co-expression network. The heatmap indicates each module's correlation with sex (red and green colours indicate high and low correlation, respectively; on a scale from -1 to 1). The corresponding *P*-value for each correlation is shown in brackets. Computations were carried out using the R package 'WGCNA'.

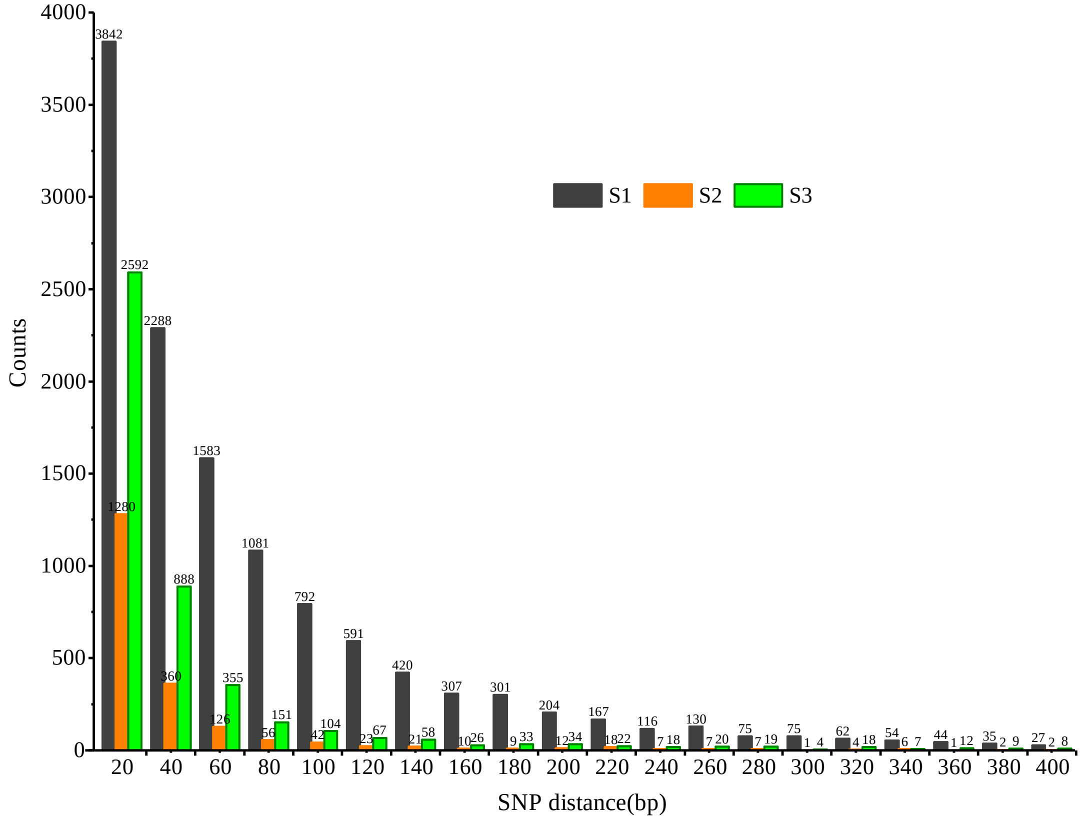


**Fig. S11. The interval of phased SNPs in the sex-determination region of gingko.** The average interval distances of the phased SNPs in S1, S2, and S3 are 83 bp, 988 bp and 303 bp, respectively.

**Table S1. Statistics of the ginkgo genome assembly[1].**

| | Scaffold (original) | Contig (original) | Scaffold (after Hi-C) | Contig (after Hi-C) |
|---|---|---|---|---|
| Total number | 169,974 | 701,988 | 76,441 | 613,806 |
| Total length (bp) | 9,626,492,610 | 9,133,153,507 | 9,554,896,939 | 9,064,524,025 |
| Average length (bp) | 56635 | 13010 | 124997 | 14767 |
| N50[2] length (bp) | 1,531,366 | 54,750 | 724,988,903 | 54,114 |
| N90[2] length (bp) | 302,221 | 10,447 | 634,527,408 | 10,334 |
| GC content (%) | 34.92 | 34.92 | 34.93 | 34.93 |

1. Sequences with a length longer than 500 bp were scored.
2. N50 length means the length for which 50% of all bases in the sequences are longer than this length. N90 length means 90% (of the total length) of the sequences are longer than this length.

**Table S2. Length of the 12 anchored ginkgo chromosomes.**

| Chromosome ID | Length (bp) |
|---|---|
| Chr1 | 1,132,477,351 |
| Chr2 | 784,418,449 |
| Chr3 | 758,283,090 |
| Chr4 | 756,296,999 |
| Chr5 | 734,512,823 |
| Chr6 | 724,988,903 |
| Chr7 | 724,829,877 |
| Chr8 | 717,075,685 |
| Chr9 | 708,856,702 |
| Chr10 | 706,912,260 |
| Chr11 | 647,862,299 |
| Chr12 | 634,527,408 |

**Table S3. Sample information and sequencing results of male and female ginkgo specimens.**

| Sample ID | Female/Male | Sequencing depth | Mapping rate |
|---|---|---|---|
| 5IL05 | F | 9.10 | 92.60 |
| ALBF1 | F | 6.80 | 94.92 |
| ALBF19 | F | 6.00 | 94.20 |
| ALBF21 | F | 6.60 | 94.25 |
| ALBF3 | F | 6.10 | 95.20 |
| ALBF8 | F | 5.70 | 95.06 |
| ATV2 | F | 9.10 | 94.50 |
| ATV3 | F | 7.80 | 94.00 |
| CQSZ01 | F | 6.60 | 93.92 |
| CQSZ03 | F | 6.50 | 95.19 |
| CQSZ05 | F | 6.30 | 94.66 |
| CQSZ07 | F | 5.70 | 95.04 |
| CQSZ11 | F | 6.00 | 95.05 |
| CQSZ14 | F | 6.60 | 93.48 |
| CQSZ15 | F | 6.40 | 94.83 |
| CQSZ16 | F | 6.70 | 94.39 |
| CQSZ17 | F | 6.40 | 94.41 |
| CX301 | F | 6.10 | 95.00 |
| CX302 | F | 5.10 | 93.80 |
| CX306 | F | 4.30 | 93.60 |
| CX307 | F | 4.40 | 90.40 |
| CX308 | F | 5.70 | 93.10 |
| CX309 | F | 4.50 | 94.10 |
| CX310 | F | 5.80 | 94.60 |
| CX312 | F | 4.40 | 94.50 |
| CX315 | F | 8.90 | 92.60 |
| DEHD3 | F | 9.00 | 94.00 |
| DF37 | F | 6.60 | 93.80 |
| DHLY5 | F | 4.30 | 92.90 |
| DHLY8 | F | 9.30 | 94.00 |
| DHLY9 | F | 8.60 | 93.30 |
| ES03 | F | 4.40 | 92.20 |
| ES05 | F | 5.10 | 89.60 |
| ES06 | F | 4.80 | 88.10 |
| ES07 | F | 5.40 | 89.60 |
| ES08 | F | 5.40 | 95.88 |
| ES10 | F | 5.50 | 91.60 |

| | | | |
|---|---|---|---|
| ES11 | F | 5.10 | 92.80 |
| ES12 | F | 4.90 | 90.60 |
| ES14 | F | 5.00 | 91.70 |
| ESBG14 | F | 4.80 | 89.50 |
| ESBG15 | F | 5.00 | 91.60 |
| ESY20 | F | 28.30 | 95.66 |
| ESY24 | F | 5.50 | 93.17 |
| ESY27 | F | 6.50 | 93.18 |
| ESY34 | F | 5.30 | 93.32 |
| ESY36 | F | 5.50 | 93.76 |
| ESY45 | F | 4.00 | 93.66 |
| ESY47 | F | 6.60 | 94.21 |
| ESY6 | F | 29.80 | 93.76 |
| FF0590 | F | 6.60 | 93.80 |
| FF0599 | F | 6.60 | 91.30 |
| FF0600 | F | 5.60 | 94.20 |
| FF0602 | F | 6.40 | 95.10 |
| FF0603 | F | 6.40 | 94.60 |
| FF0605 | F | 6.70 | 92.80 |
| FF0608 | F | 6.60 | 94.60 |
| FF0611 | F | 6.10 | 93.20 |
| FF0615 | F | 6.30 | 94.20 |
| FF0616 | F | 6.70 | 91.80 |
| FJPC07 | F | 6.40 | 94.86 |
| FJPC19 | F | 4.10 | 94.27 |
| FJPC23 | F | 6.40 | 95.77 |
| FJPC26 | F | 3.00 | 94.31 |
| FJPC32 | F | 4.90 | 93.02 |
| FJPC33 | F | 5.50 | 95.11 |
| FJSC08 | F | 5.40 | 94.90 |
| FJSC13 | F | 5.00 | 93.47 |
| FJSC16 | F | 3.80 | 92.18 |
| FJYX52 | F | 6.40 | 95.12 |
| FJYX64 | F | 6.50 | 94.52 |
| FJYX75 | F | 6.50 | 94.37 |
| FJZH01 | F | 6.40 | 94.50 |
| FJZH03 | F | 6.50 | 94.94 |
| FJZH12 | F | 6.40 | 95.28 |
| FJZH13 | F | 6.10 | 94.95 |
| FJZH41 | F | 4.30 | 93.41 |

| | | | |
|---|---|---|---|
| FREU12 | F | 4.50 | 92.40 |
| GXLC02 | F | 6.80 | 95.30 |
| GXLC06 | F | 6.20 | 94.98 |
| GXLC11 | F | 6.60 | 94.55 |
| GXLC12 | F | 6.20 | 95.01 |
| GXLC13 | F | 6.30 | 95.09 |
| GXLC14 | F | 6.00 | 95.27 |
| GXLC15 | F | 6.10 | 95.10 |
| GXLC16 | F | 6.90 | 95.64 |
| GXLC17 | F | 6.80 | 95.16 |
| GXLC18 | F | 6.10 | 94.98 |
| GXXA02 | F | 28.40 | 96.49 |
| GXXA06 | F | 6.50 | 95.11 |
| GXXA09 | F | 6.40 | 94.82 |
| GXXA15 | F | 6.30 | 95.90 |
| GXXA24 | F | 5.60 | 95.00 |
| GXXA26 | F | 6.00 | 94.64 |
| GXXA30 | F | 5.40 | 94.83 |
| GXXA31 | F | 6.00 | 95.35 |
| GXXA34 | F | 5.70 | 95.21 |
| GXXA35 | F | 6.60 | 95.20 |
| GYSP01 | F | 6.90 | 92.00 |
| GYSP04 | F | 6.50 | 89.60 |
| GYSP05 | F | 7.20 | 94.00 |
| GYSP13 | F | 5.00 | 92.30 |
| HESX10 | F | 6.60 | 94.80 |
| HESX12 | F | 6.60 | 94.47 |
| HESX15 | F | 6.70 | 94.23 |
| HEXX1 | F | 6.30 | 94.40 |
| HEXX13 | F | 6.10 | 95.03 |
| HEXX15 | F | 6.70 | 94.23 |
| HEXX2 | F | 6.10 | 94.95 |
| HEXX7 | F | 6.30 | 94.77 |
| HEXX8 | F | 6.40 | 94.61 |
| HP02 | F | 12.10 | 92.50 |
| HP03 | F | 9.40 | 91.20 |
| HP04 | F | 9.30 | 89.50 |
| HP05 | F | 4.20 | 94.00 |
| HP07 | F | 7.40 | 90.70 |
| HP08 | F | 5.00 | 92.10 |

| | | | |
|---|---|---|---|
| HX01 | F | 8.70 | 88.00 |
| HX04 | F | 4.60 | 84.40 |
| HX09 | F | 4.30 | 82.50 |
| HX11 | F | 4.20 | 93.40 |
| HX13 | F | 8.80 | 91.80 |
| HX19 | F | 5.00 | 92.90 |
| HX22 | F | 7.70 | 91.60 |
| HX28 | F | 5.30 | 92.70 |
| HX29 | F | 4.80 | 93.50 |
| HX30 | F | 4.50 | 93.40 |
| HX33 | F | 5.30 | 93.50 |
| HX35 | F | 5.00 | 91.50 |
| HX36 | F | 4.80 | 91.00 |
| HX38 | F | 4.90 | 93.70 |
| HX39 | F | 4.70 | 93.80 |
| JF304 | F | 7.00 | 89.90 |
| JF312 | F | 7.80 | 92.50 |
| JF314 | F | 10.00 | 92.70 |
| JFDL48 | F | 5.80 | 94.05 |
| JFDL50 | F | 6.40 | 93.53 |
| JFDY01 | F | 6.50 | 93.56 |
| JFDY02 | F | 6.60 | 94.37 |
| JFDY03 | F | 6.50 | 93.58 |
| JFDY04 | F | 6.20 | 91.83 |
| JFDY05 | F | 6.60 | 94.11 |
| JFDY06 | F | 6.80 | 91.13 |
| JFDY07 | F | 6.70 | 95.40 |
| JFPY20 | F | 7.30 | 90.60 |
| JFPY21 | F | 5.10 | 94.20 |
| JR03 | F | 4.20 | 93.30 |
| JSYX06 | F | 4.70 | 89.50 |
| JSYX08 | F | 4.70 | 90.60 |
| JZ01 | F | 4.20 | 92.70 |
| JZ02 | F | 5.00 | 93.90 |
| JZ06 | F | 4.40 | 92.20 |
| JZ07 | F | 4.40 | 93.70 |
| JZ09 | F | 10.50 | 94.30 |
| JZ14 | F | 7.10 | 92.50 |
| JZ15 | F | 9.70 | 92.70 |
| KX1 | F | 6.60 | 94.76 |

| | | | |
|---|---|---|---|
| KX6 | F | 6.50 | 94.78 |
| LYG06 | F | 5.50 | 93.60 |
| LYG10 | F | 10.30 | 94.60 |
| LYG12 | F | 12.10 | 94.20 |
| LYG15 | F | 4.80 | 93.20 |
| MAPH4 | F | 5.60 | 93.80 |
| MT08 | F | 6.50 | 94.20 |
| NC07 | F | 4.80 | 94.40 |
| NC08 | F | 7.00 | 94.90 |
| NC09 | F | 5.10 | 86.30 |
| NX01 | F | 6.20 | 95.40 |
| NX05 | F | 6.20 | 95.23 |
| NX11 | F | 6.00 | 95.51 |
| NX20 | F | 6.40 | 95.02 |
| NX25 | F | 6.40 | 95.48 |
| NX34 | F | 6.20 | 95.59 |
| NX43 | F | 6.50 | 95.49 |
| NX46 | F | 6.30 | 95.75 |
| PAPH2 | F | 4.60 | 93.20 |
| PXLM15 | F | 3.70 | 92.50 |
| RZ12 | F | 4.20 | 94.00 |
| RZ15 | F | 5.00 | 92.60 |
| SCQL06 | F | 6.30 | 95.04 |
| SCQL08 | F | 6.40 | 95.55 |
| SCWY01 | F | 6.50 | 94.77 |
| SCWY03 | F | 6.60 | 95.55 |
| SG07 | F | 4.40 | 94.70 |
| SKK355 | F | 6.40 | 93.30 |
| TM010 | F | 8.20 | 93.00 |
| TM01L | F | 5.90 | 89.50 |
| TM02L | F | 7.90 | 91.80 |
| TM038 | F | 3.90 | 93.90 |
| TM03L | F | 7.60 | 91.50 |
| TM049 | F | 6.30 | 92.60 |
| TM04L | F | 4.70 | 91.40 |
| TM052 | F | 8.80 | 92.50 |
| TM05L | F | 5.40 | 94.00 |
| TM064 | F | 7.40 | 91.50 |
| TM065 | F | 8.20 | 94.20 |
| TM067 | F | 8.80 | 93.30 |

| | | | |
|---|---|---|---|
| TM068 | F | 8.90 | 94.00 |
| TM070 | F | 8.90 | 93.96 |
| TM106 | F | 30.20 | 96.10 |
| TM142 | F | 4.30 | 88.10 |
| TM147 | F | 8.60 | 93.70 |
| TM169 | F | 30.60 | 95.75 |
| TM191 | F | 6.10 | 92.10 |
| TM226 | F | 8.10 | 94.60 |
| TM289 | F | 6.30 | 93.00 |
| TM358 | F | 9.00 | 93.40 |
| TM407 | F | 5.90 | 91.50 |
| TM425 | F | 3.70 | 98.10 |
| TM428 | F | 4.40 | 98.10 |
| TM432 | F | 3.90 | 97.40 |
| TM439 | F | 4.60 | 97.90 |
| TM474 | F | 7.80 | 95.00 |
| TMAB0715 | F | 4.50 | 98.00 |
| TMAB0722 | F | 2.90 | 97.90 |
| WA01 | F | 6.40 | 94.58 |
| WA08 | F | 6.20 | 95.55 |
| WA11 | F | 6.40 | 95.08 |
| WA13 | F | 6.40 | 95.64 |
| WA22 | F | 6.50 | 94.92 |
| WA24 | F | 5.20 | 95.40 |
| WA25 | F | 5.30 | 95.79 |
| WA27 | F | 5.80 | 95.77 |
| WC006 | F | 5.80 | 94.50 |
| WC012 | F | 6.30 | 94.10 |
| WC027 | F | 6.40 | 92.70 |
| WC034 | F | 6.50 | 93.80 |
| WC040 | F | 6.70 | 94.60 |
| WC047 | F | 6.50 | 94.60 |
| WC054 | F | 6.70 | 91.30 |
| WC073 | F | 6.40 | 93.80 |
| WC092 | F | 6.20 | 94.30 |
| WC101 | F | 6.20 | 95.20 |
| WC103 | F | 6.40 | 94.60 |
| WCBY206 | F | 4.20 | 92.80 |
| WCNT52 | F | 4.40 | 88.20 |
| WCNT81 | F | 4.60 | 91.80 |

| | | | |
|---|---|---|---|
| WCSE25 | F | 4.60 | 91.80 |
| WCSE31 | F | 8.00 | 92.30 |
| WCSE35 | F | 4.90 | 93.50 |
| WCSE39 | F | 4.60 | 88.10 |
| WCSE71 | F | 5.30 | 84.50 |
| WX1 | F | 6.70 | 94.94 |
| WX10 | F | 6.10 | 94.76 |
| WX4 | F | 6.50 | 94.25 |
| WX5 | F | 6.50 | 94.46 |
| WX6 | F | 6.90 | 94.96 |
| WX7 | F | 6.90 | 93.86 |
| WX8 | F | 1.50 | 94.15 |
| XC04 | F | 4.60 | 92.90 |
| XX05 | F | 5.30 | 93.00 |
| XX06 | F | 8.50 | 94.70 |
| XX08 | F | 8.20 | 94.60 |
| XX13 | F | 4.30 | 93.40 |
| XX14 | F | 4.60 | 92.90 |
| YNTC17 | F | 6.40 | 95.44 |
| YNTC19 | F | 6.30 | 94.61 |
| YNTC22 | F | 6.20 | 94.72 |
| YNTC25 | F | 6.60 | 94.44 |
| YNTC26 | F | 6.20 | 94.67 |
| YNTC27 | F | 5.90 | 94.27 |
| YX08 | F | 4.70 | 93.00 |
| YX09 | F | 8.80 | 93.30 |
| YX10 | F | 4.80 | 92.80 |
| ZH04 | F | 7.50 | 85.50 |
| ALBF41 | M | 6.60 | 95.28 |
| ATV1 | M | 9.40 | 94.90 |
| ATV4 | M | 10.60 | 93.38 |
| CC22 | M | 4.40 | 94.20 |
| CQSZ08 | M | 5.30 | 94.24 |
| CXS012 | M | 9.20 | 93.00 |
| CXS019 | M | 8.40 | 92.70 |
| DEW2 | M | 4.60 | 94.20 |
| ESBG13 | M | 4.20 | 90.60 |
| ESY30 | M | 6.80 | 93.40 |
| FJPC20 | M | 5.70 | 94.31 |
| FJPC29 | M | 6.30 | 94.25 |

| | | | |
|---|---|---|---|
| FJSC04 | M | 5.30 | 94.23 |
| FJSC07 | M | 5.10 | 93.93 |
| FJSC30 | M | 6.10 | 93.66 |
| FJYX22 | M | 6.30 | 95.50 |
| FJYX24 | M | 6.20 | 93.70 |
| FJYX51 | M | 6.60 | 94.98 |
| FJYX71 | M | 6.60 | 95.23 |
| FJYX79 | M | 6.50 | 94.90 |
| FJZH09 | M | 6.60 | 94.78 |
| FJZH14 | M | 6.30 | 93.75 |
| FJZH19 | M | 6.60 | 94.98 |
| FJZH26 | M | 6.40 | 93.00 |
| FJZH27 | M | 6.20 | 95.09 |
| FREU14 | M | 4.30 | 94.60 |
| FRFP1 | M | 8.90 | 93.90 |
| FUK04 | M | 9.30 | 93.10 |
| FUK06 | M | 7.60 | 93.60 |
| FUK15 | M | 5.60 | 92.20 |
| FUK16 | M | 5.40 | 92.60 |
| FUK17 | M | 4.90 | 92.50 |
| GYSP06 | M | 7.10 | 92.20 |
| HESX1 | M | 29.20 | 96.38 |
| HESX16 | M | 5.90 | 94.25 |
| HESX19 | M | 5.60 | 94.82 |
| HESX20 | M | 6.40 | 95.22 |
| HESX4 | M | 6.10 | 94.56 |
| HESX6 | M | 5.90 | 94.13 |
| HESX8 | M | 6.60 | 94.73 |
| HEXX10 | M | 6.00 | 95.15 |
| HEXX12 | M | 6.10 | 94.75 |
| HEXX18 | M | 6.70 | 93.65 |
| HEXX4 | M | 6.10 | 94.56 |
| HP06 | M | 8.70 | 93.40 |
| HP09 | M | 8.30 | 93.30 |
| HX23 | M | 4.90 | 91.10 |
| HX32 | M | 4.30 | 93.30 |
| JFDL33 | M | 29.60 | 95.75 |
| JFDL39 | M | 27.90 | 96.31 |
| JFPY16 | M | 5.10 | 93.90 |
| JFPY19 | M | 8.20 | 92.50 |

| | | | |
|---|---|---|---|
| JR07 | M | 4.20 | 94.00 |
| JZ04 | M | 4.30 | 93.00 |
| JZ11 | M | 9.00 | 91.40 |
| KX13 | M | 6.80 | 94.60 |
| KX14 | M | 6.40 | 93.77 |
| KX19 | M | 6.50 | 94.62 |
| KX34 | M | 6.60 | 93.67 |
| KX41 | M | 6.40 | 93.48 |
| KX42 | M | 6.80 | 94.19 |
| KYS371 | M | 6.70 | 93.60 |
| MAPH3 | M | 5.40 | 92.40 |
| MAPH5 | M | 8.50 | 92.00 |
| NLEU1 | M | 5.90 | 94.70 |
| NX31 | M | 6.10 | 95.69 |
| NX39 | M | 6.20 | 95.56 |
| NY01 | M | 4.40 | 93.70 |
| PAPH1 | M | 4.40 | 92.60 |
| QY08 | M | 4.30 | 93.10 |
| RZ08 | M | 5.00 | 92.90 |
| SCQL01 | M | 6.10 | 93.51 |
| SCQL02 | M | 6.20 | 93.50 |
| SCQL03 | M | 6.30 | 94.06 |
| SCQL04 | M | 6.30 | 94.71 |
| SCWY04 | M | 6.40 | 95.43 |
| SCWY06 | M | 6.40 | 95.44 |
| SCWY07 | M | 6.50 | 95.53 |
| SCWY08 | M | 6.80 | 95.26 |
| SCWY12 | M | 5.60 | 94.92 |
| SCWY15 | M | 5.10 | 94.04 |
| SCWY23 | M | 6.60 | 93.36 |
| TM054 | M | 3.10 | 97.60 |
| TM140 | M | 3.50 | 96.90 |
| TM171 | M | 24.90 | 95.55 |
| TM179 | M | 8.50 | 93.00 |
| TM190 | M | 7.20 | 90.40 |
| TM218 | M | 7.80 | 93.50 |
| TM270 | M | 6.50 | 90.60 |
| TMAB0759-2 | M | 5.30 | 90.70 |
| WA07 | M | 6.40 | 94.67 |
| WCBL5 | M | 4.50 | 89.90 |

| | | | |
|---|---|---|---|
| WCSE57 | M | 4.60 | 93.00 |
| WCSE70 | M | 5.00 | 90.70 |
| WX9 | M | 6.20 | 93.89 |
| YX03 | M | 7.90 | 91.30 |
| ZH02 | M | 9.20 | 87.50 |

**Table S4. Divergence time (MYA) of S1 and S2 between 4 males and 5 females.**

| S1 | Male1 | Male2 | Male3 | Male4 | Female1 | Female1 | Female3 | Female4 | Female5 |
|---|---|---|---|---|---|---|---|---|---|
| Male1 | 0.00 | 0.01 | 2.76 | 5.37 | 13.73 | 13.88 | 13.88 | 15.07 | 15.30 |
| Male2 | 2.91 | 0.00 | 1.27 | 5.67 | 15.30 | 15.37 | 15.45 | 16.57 | 16.19 |
| Male3 | 2.76 | 0.01 | 0.00 | 5.52 | 15.15 | 15.22 | 15.30 | 16.42 | 16.04 |
| Male4 | 5.37 | 0.03 | 5.52 | 0.00 | 11.04 | 11.12 | 11.19 | 12.31 | 12.09 |
| Female1 | 13.73 | 0.07 | 15.15 | 11.04 | 0.00 | 0.22 | 0.30 | 1.72 | 1.94 |
| Female2 | 13.88 | 0.07 | 15.22 | 11.12 | 0.22 | 0.00 | 0.37 | 1.79 | 2.01 |
| Female3 | 13.88 | 0.07 | 15.30 | 11.19 | 0.30 | 0.37 | 0.00 | 1.79 | 1.94 |
| Female4 | 15.07 | 0.07 | 16.42 | 12.31 | 1.72 | 1.79 | 1.79 | 0.00 | 0.90 |
| Female5 | 15.30 | 0.07 | 16.04 | 12.09 | 1.94 | 2.01 | 1.94 | 0.90 | 0.00 |

| S3 | Male1 | Male2 | Male3 | Male4 | Female1 | Female1 | Female3 | Female4 | Female5 |
|---|---|---|---|---|---|---|---|---|---|
| Male1 | 0.00 | 0.02 | 4.10 | 7.69 | 9.93 | 11.19 | 9.85 | 11.27 | 11.19 |
| Male2 | 3.96 | 0.00 | 3.73 | 6.94 | 10.97 | 10.60 | 11.12 | 10.60 | 10.60 |
| Male3 | 4.10 | 0.02 | 0.00 | 7.16 | 11.34 | 10.90 | 11.34 | 11.12 | 11.12 |
| Male4 | 7.69 | 0.03 | 7.16 | 0.00 | 5.60 | 4.48 | 5.45 | 4.93 | 4.93 |
| Female1 | 9.93 | 0.05 | 11.34 | 5.60 | 0.00 | 1.49 | 0.52 | 1.57 | 1.57 |
| Female2 | 11.19 | 0.05 | 10.90 | 4.48 | 1.49 | 0.00 | 1.72 | 0.75 | 0.75 |
| Female3 | 9.85 | 0.05 | 11.34 | 5.45 | 0.52 | 1.72 | 0.00 | 1.72 | 1.72 |
| Female4 | 11.27 | 0.05 | 11.12 | 4.93 | 1.57 | 0.75 | 1.72 | 0.00 | 0.15 |
| Female5 | 11.19 | 0.05 | 11.12 | 4.93 | 1.57 | 0.75 | 1.72 | 0.15 | 0.00 |

**Table S5. Genes located in the sex-determination region of ginkgo.**

| Gene ID | Homologous protein[1] | Annotation |
|---|---|---|
| ***Gb_15883*** | RR12 | Response regulator 12; Transcriptional activator that binds specifically to the DNA sequence 5'-[AG]GATT-3'. Functions as a response regulator involved in His-to-Asp phosphorelay signal transduction system. Could directly activate some type-A response regulators in response to cytokinin. A Y-Encoded Suppressor of Feminization Arose via Lineage-Specific Duplication of a Cytokinin Response Regulator in Kiwifruit. |
| ***Gb_15884*** | RR2 | Response regulator 2; Transcriptional activator that binds specifically to the DNA sequence 5'-[AG]GATT-3'. Functions as a response regulator involved in His-to-Asp phosphorelay signal transduction system. Could directly activate some type-A response regulators in response to cytokinins. A Y-Encoded Suppressor of Feminization Arose via Lineage-Specific Duplication of a Cytokinin Response Regulator in Kiwifruit. |
| ***Gb_15885*** | AT5G46910 | Transcription factor jumonji and C5HC2 type zinc finger domain-containing protein |
| | REF6 | Relative of early flowering 6; Histone demethylase that demethylates 'Lys-27' (H3K27me) of histone H3. Demethylates both tri- (H3K27me3) and di-methylated (H3K27me2) H3K27me. Demethylates also H3K4me3/2 and H3K36me3/2 in an in vitro assay. Involved in the transcriptional regulation of hundreds of genes regulating developmental patterning and responses to various stimuli. Involved in the regulation of flowering time by repressing FLOWERING LOCUS C (FLC) expression |
| | MEE27 | Maternal effect embryo arrest 27; Histone demethylase that demethylates 'Lys-4' (H3K4me) of histone H3 with a specific activity for H3K4me3. No activity on H3K4me2, H3K4me1, H3K9me3/2, H3K27me3/2 and H3K36me3/2. Involved in the control of flowering time by demethylating H3K4me3 at the FLC locus and repressing its expression. The repression of FLC level and reduction in H3K4me3 at the FLC locus results in induction of the flowering activator FT, which is a downstream target of FLC |
| | JMJ18 | Jumonji domain-containing protein 18; Histone demethylase that demethylates 'Lys-4' (H3K4me) of histone H3 with a specific activity for H3K4me3 and H3K4me2. No activity on H3K9me3/2, H3K27me3/2 and H3K36me3/2. Involved in the control of flowering time by demethylating H3K4me3 at the FLC locus and |

| | | |
|---|---|---|
| | | repressing its expression. The repression of FLC level and reduction in H3K4me3 at the FLC locus results in induction of the flowering activator FT, which is a downstream target of FLC |
| | ELF6 | EARLY FLOWERING 6; Acts probably as a histone 'Lys-4' (H3K4me) demethylase. Involved in transcriptional gene regulation. Acts as a repressor of the photoperiodic flowering pathway and of FT. Binds around the transcription start site of the FT locus |
| ***Gb_15886*** | AT4G31910 | BR-related AcylTransferase1, Brassinosteroid control of sex determination in maize |
| | RWP1 | REDUCED LEVELS OF WALL-BOUND PHENOLICS 1; Involved in the synthesis of aromatics of the suberin polymer. Specifically affects the accumulation of the ferulate constituent of suberin in roots and seeds, but has no effect on the content of p-coumarate or sinapate |
| ***Gb_28584*** | AT4G04480 | Uncharacterized protein |
| | AT4G22030 | Putative F-box protein |
| ***Gb_28585*** | - - | [STRING found no matching protein in its database] |
| ***Gb_28586*** | SLO2 | SLOW GROWTH 2 |
| | AT5G16860 | Pentatricopeptide repeat-containing protein |
| | DOT4 | DEFECTIVELY ORGANIZED TRIBUTARIES 4; Plays a major role in single RNA editing events in chloroplasts. Acts as a site-recognition transacting factor involved in the edition of the unique site (corresponding to cytidine-488) of rpoC1, which is a plastid-encoded subunit of the chloroplast DNA-directed RNA polymerase. May provide the catalytic activity for editing site conversion (PubMed:24194514). Involved in leaf vasculature patterning (PubMed:18643975) |
| | OTP84 | ORGANELLE TRANSCRIPT PROCESSING 84; Involved in RNA editing events in chloroplasts. Required for the editing of a single site in ndhB and ndhF transcripts, which are two plastid-encoded subunits of the chloroplast NAD(P)H dehydrogenase (NDH) complex. Required for the editing of a single site in psbZ. Required for optimal activity of the NDH complex of the photosynthetic electron transport chain |
| | AT2G22070 | Pentatricopeptide repeat-containing protein |
| ***Gb_28587*** | AGL6 | AGAMOUS-like 6; Probable transcription factor. Forms a heterodimer via the K-box domain with AG, that could be involved in genes regulation during floral meristem development |
| | AGL13 | AGAMOUS-like 13; Probable transcription factor |

| | | |
|---|---|---|
| | AGL8 | AGAMOUS-like 8; Probable transcription factor that promotes early floral meristem identity in synergy with APETALA1 and CAULIFLOWER. Is required subsequently for the transition of an inflorescence meristem into a floral meristem. Seems to be partially redundant to the function of APETALA1 and CAULIFLOWER in the up-regulation of LEAFY. Is also required for normal pattern of cell division, expansion and differentiation during morphogenesis of the silique. Probably not required for fruit elongation but instead is required to prevent ectopic activity of IND |
| | AG | AGAMOUS; Probable transcription factor involved in the control of organ identity during the early development of flowers. Is required for normal development of stamens and carpels in the wild-type flower. Plays a role in maintaining the determinacy of the floral meristem. Acts as C class cadastral protein by repressing the A class floral homeotic genes like APETALA1. Forms a heterodimer via the K-box domain with either SEPALATTA1/AGL2, SEPALATTA2/AGL4, SEPALLATA3/AGL9 or AGL6 that could be involved in genes regulation during floral meristem development. Controls AHL21/GIK, a multifunct [...] |
| | PI | PISTILLATA; Probable transcription factor involved in the genetic control of flower development. Is required for normal development of petals and stamens in the wild-type flower. Forms a heterodimer with APETALA3 that is required for autoregulation of both AP3 and PI genes. AP3/PI heterodimer interacts with APETALA1 or SEPALLATA3 to form a ternary complex that could be responsible for the regulation of the genes involved in the flower development. AP3/PI heterodimer activates the expression of NAP |
| ***Gb_28588*** | AT3G01930 | Major facilitator protein |
| | AT5G14120 | Major facilitator protein |
| ***Gb_28589*** | AT4G31020 | Esterase/lipase domain-containing protein |
| | AT2G24320 | alpha/beta-Hydrolases superfamily protein |
| | AT3G30380 | Esterase/lipase domain-containing protein |
| | AT5G38220 | Hydrolase |
| | AT4G24760 | Esterase/lipase domain-containing protein |
| ***Gb_30340*** | - - | [STRING found no matching protein in its database] |
| ***Gb_30341*** | AT3G17450 | hAT dimerisation domain-containing protein |
| ***Gb_30342*** | AT3G17450 | hAT dimerisation domain-containing protein |
| | AT1G79740 | hAT transposon superfamily |

| | | |
|---|---|---|
| ***Gb_30343*** | BIO1 | Biotin auxotroph 1; Bifunctional enzyme that catalyzes two different reactions involved in the biotin biosynthesis |
| ***Gb_30344*** | AT4G30310 | FGGY family of carbohydrate kinase |
| ***Gb_30345*** | AT3G17450 | hAT dimerisation domain-containing protein |
| | AT3G13030 | hAT transposon superfamily protein |

1. Gene functional annotation was conducted by searching for the homologous proteins in the STIRING protein database. The homologous proteins were indicated.

**Table S6. Overview of ginkgo transcriptome data.**

| No. | Sample | Sex | Tissue | Clean reads | Clean base |
|---|---|---|---|---|---|
| 1 | F1-7A | Female | Cones | 76,714,484 | 6,520,731,140 |
| 2 | F1-7A | Female | Cones | 66,016,380 | 5,611,392,300 |
| 3 | F2-4A | Female | Cones | 71,077,868 | 6,041,618,780 |
| 4 | F2-6A | Female | Cones | 81,984,378 | 6,968,672,130 |
| 5 | F4-4A | Female | Cones | 114,362,702 | 9,720,829,670 |
| 6 | F4-6A | Female | Cones | 76,515,508 | 6,503,818,180 |
| 7 | F4-7A | Female | Cones | 86,586,060 | 7,359,815,100 |
| 8 | LF1-7A | Female | Leaves | 83,875,480 | 7,129,415,800 |
| 9 | LF1-7A | Female | Leaves | 86,618,294 | 7,362,554,990 |
| 10 | LF2-4A | Female | Leaves | 80,750,060 | 6,863,755,100 |
| 11 | LF2-6A | Female | Leaves | 70,310,138 | 5,976,361,730 |
| 12 | LF4-4A | Female | Leaves | 60,373,628 | 5,131,758,380 |
| 13 | LF4-6A | Female | Leaves | 50,631,002 | 4,303,635,170 |
| 14 | LF4-7A | Female | Leaves | 100,929,592 | 8,579,015,320 |
| 15 | M1-1A | Male | Cones | 91,877,286 | 7,809,569,310 |
| 16 | M1-2A | Male | Cones | 56,926,514 | 4,838,753,690 |
| 17 | M1-5A | Male | Cones | 77,559,492 | 6,592,556,820 |
| 18 | M2-1A | Male | Cones | 74,288,854 | 6,314,552,590 |
| 19 | M2-2A | Male | Cones | 83,395,228 | 7,088,594,380 |
| 20 | M2-5A | Male | Cones | 66,649,044 | 5,665,168,740 |
| 21 | M3-1A | Male | Cones | 77,078,570 | 6,551,678,450 |
| 22 | M3-2A | Male | Cones | 78,776,948 | 6,696,040,580 |
| 23 | M3-5A | Male | Cones | 79,665,954 | 5,576,616,780 |
| 24 | LM1-1A | Male | Leaves | 95,422,690 | 8,110,928,650 |
| 25 | LM1-2A | Male | Leaves | 85,128,118 | 7,235,890,030 |
| 26 | LM1-5A | Male | Leaves | 77,921,570 | 6,623,333,450 |
| 27 | LM2-1A | Male | Leaves | 86,257,388 | 7,331,877,980 |
| 28 | LM2-2A | Male | Leaves | 69,429,484 | 5,901,506,140 |
| 29 | LM2-5A | Male | Leaves | 73,534,238 | 6,250,410,230 |
| 30 | LM3-1A | Male | Leaves | 86,996,506 | 7,394,703,010 |
| 31 | LM3-2A | Male | Leaves | 92,593,420 | 7,870,440,700 |
| 32 | LM3-5A | Male | Leaves | 70,465,246 | 5,989,545,910 |

**Table S7. Specifically expressed genes (SEGs) in either male or female ginkgo cones. TPM denotes transcripts per million; f, female; m, male; SD, standard deviation.**

| Gene | 7 females avg (TPM) | 7 females SD (TPM) | 9 males avg (TPM) | 9 males SD (TPM) | Log (F/M) |
|---|---|---|---|---|---|
| *Gb_36895* | 0.55 | 0.20 | 4.66 | 2.34 | -0.32 |
| *Gb_26014* | 0.50 | 0.23 | 4.55 | 2.54 | -0.31 |
| *Gb_25620* | 0.55 | 0.27 | 5.31 | 2.61 | -0.31 |
| *Gb_15682* | 0.51 | 0.18 | 5.53 | 3.46 | -0.29 |
| *Gb_17880* | 0.65 | 0.29 | 8.44 | 3.80 | -0.27 |
| *Gb_39464* | 0.62 | 0.25 | 10.50 | 4.68 | -0.24 |
| *Gb_35206* | 0.42 | 0.17 | 8.76 | 5.22 | -0.23 |
| *Gb_01351* | 0.45 | 0.18 | 9.47 | 4.96 | -0.23 |
| *Gb_14146* | 0.31 | 0.22 | 6.66 | 4.28 | -0.23 |
| *Gb_04497* | 0.36 | 0.16 | 7.58 | 5.71 | -0.23 |
| *Gb_22052* | 0.36 | 0.16 | 7.58 | 5.71 | -0.23 |
| *Gb_39730* | 0.40 | 0.14 | 10.98 | 8.00 | -0.21 |
| *Gb_26320* | 0.31 | 0.13 | 8.56 | 4.29 | -0.21 |
| *Gb_40846* | 0.44 | 0.35 | 12.77 | 6.89 | -0.21 |
| *Gb_33168* | 0.42 | 0.24 | 12.20 | 6.90 | -0.21 |
| *Gb_01318* | 0.32 | 0.24 | 10.64 | 9.14 | -0.20 |
| *Gb_23879* | 0.24 | 0.10 | 8.47 | 6.95 | -0.20 |
| *Gb_29151* | 0.36 | 0.29 | 13.71 | 14.21 | -0.19 |
| *Gb_12049* | 0.21 | 0.13 | 8.14 | 2.31 | -0.19 |
| *Gb_39463* | 0.25 | 0.35 | 9.97 | 4.77 | -0.19 |
| *Gb_38997* | 0.71 | 0.14 | 28.08 | 18.29 | -0.19 |
| *Gb_11052* | 0.20 | 0.09 | 8.67 | 5.03 | -0.18 |
| *Gb_20473* | 0.38 | 0.19 | 20.25 | 15.32 | -0.17 |
| *Gb_25546* | 0.42 | 0.12 | 22.82 | 13.76 | -0.17 |
| *Gb_38572* | 0.44 | 0.31 | 25.56 | 18.30 | -0.17 |
| *Gb_08061* | 0.32 | 0.17 | 18.94 | 20.14 | -0.17 |
| *Gb_13578* | 0.23 | 0.13 | 13.61 | 15.61 | -0.17 |
| *Gb_21779* | 0.23 | 0.13 | 13.61 | 15.61 | -0.17 |
| *Gb_00022* | 0.23 | 0.13 | 13.61 | 15.61 | -0.17 |
| *Gb_32151* | 0.46 | 0.22 | 33.78 | 31.61 | -0.16 |
| *Gb_20316* | 0.18 | 0.25 | 13.99 | 11.90 | -0.16 |
| *Gb_41020* | 0.31 | 0.14 | 27.55 | 15.71 | -0.15 |
| *Gb_02087* | 0.21 | 0.11 | 19.21 | 9.95 | -0.15 |
| *Gb_30968* | 0.19 | 0.13 | 19.56 | 6.75 | -0.15 |
| *Gb_03601* | 0.17 | 0.22 | 20.53 | 13.70 | -0.14 |
| *Gb_34102* | 0.10 | 0.05 | 12.78 | 7.21 | -0.14 |
| *Gb_04228* | 0.62 | 0.20 | 80.54 | 59.10 | -0.14 |
| *Gb_02979* | 0.34 | 0.27 | 51.26 | 64.62 | -0.14 |
| *Gb_01997* | 0.37 | 0.21 | 69.85 | 57.35 | -0.13 |

| | | | | | |
|---|---|---|---|---|---|
| *Gb_26627* | 0.53 | 0.42 | 101.65 | 50.59 | -0.13 |
| *Gb_19431* | 0.36 | 0.21 | 72.67 | 68.14 | -0.13 |
| *Gb_07514* | 0.26 | 0.35 | 53.34 | 59.78 | -0.13 |
| *Gb_19318* | 0.09 | 0.08 | 18.86 | 8.93 | -0.13 |
| *Gb_13191* | 0.12 | 0.14 | 26.43 | 33.67 | -0.13 |
| *Gb_28689* | 0.25 | 0.18 | 57.97 | 49.48 | -0.13 |
| *Gb_39812* | 0.28 | 0.35 | 66.72 | 106.18 | -0.13 |
| *Gb_22366* | 0.16 | 0.30 | 54.92 | 92.49 | -0.12 |
| *Gb_26208* | 0.09 | 0.10 | 31.49 | 35.53 | -0.12 |
| *Gb_17168* | 0.04 | 0.07 | 15.14 | 10.03 | -0.12 |
| *Gb_07869* | 0.02 | 0.03 | 11.14 | 10.09 | -0.11 |
| *Gb_00660* | 0.30 | 0.14 | 157.84 | 128.68 | -0.11 |
| *Gb_07624* | 0.10 | 0.05 | 56.42 | 50.69 | -0.11 |
| *Gb_14191* | 0.19 | 0.13 | 115.38 | 51.76 | -0.11 |
| *Gb_18506* | 0.05 | 0.08 | 31.26 | 29.86 | -0.11 |
| *Gb_36229* | 0.06 | 0.03 | 38.59 | 47.75 | -0.11 |
| *Gb_08982* | 0.31 | 0.24 | 231.43 | 365.80 | -0.10 |
| *Gb_41246* | 0.08 | 0.13 | 71.46 | 58.36 | -0.10 |
| *Gb_40006* | 0.03 | 0.06 | 26.11 | 32.31 | -0.10 |
| *Gb_25367* | 0.02 | 0.03 | 20.19 | 14.77 | -0.10 |
| *Gb_27840* | 0.21 | 0.17 | 233.75 | 205.11 | -0.10 |
| *Gb_34493* | 0.03 | 0.05 | 33.53 | 24.71 | -0.10 |
| *Gb_13289* | 0.10 | 0.16 | 124.23 | 128.21 | -0.10 |
| *Gb_13127* | 0.09 | 0.24 | 124.51 | 113.36 | -0.10 |
| *Gb_24686* | 0.12 | 0.19 | 166.83 | 107.65 | -0.10 |
| *Gb_11266* | 0.12 | 0.25 | 166.04 | 247.92 | -0.10 |
| *Gb_09536* | 0.12 | 0.16 | 178.12 | 199.58 | -0.10 |
| *Gb_05378* | 0.02 | 0.02 | 23.39 | 15.10 | -0.10 |
| *Gb_37318* | 0.17 | 0.14 | 247.08 | 208.75 | -0.09 |
| *Gb_02049* | 0.02 | 0.03 | 36.84 | 43.48 | -0.09 |
| *Gb_32354* | 0.02 | 0.04 | 50.29 | 42.98 | -0.09 |
| *Gb_40539* | 0.03 | 0.03 | 97.77 | 89.04 | -0.09 |
| *Gb_31417* | 0.03 | 0.07 | 92.60 | 38.83 | -0.09 |
| *Gb_19469* | 0.13 | 0.28 | 471.53 | 424.75 | -0.08 |
| *Gb_10500* | 0.04 | 0.06 | 213.74 | 134.09 | -0.08 |
| *Gb_23376* | 0.03 | 0.04 | 218.62 | 190.98 | -0.08 |
| *Gb_32463* | 0.16 | 0.20 | 996.16 | 837.42 | -0.08 |
| *Gb_23369* | 0.11 | 0.23 | 691.24 | 674.68 | -0.08 |
| *Gb_38687* | 0.06 | 0.15 | 395.58 | 279.12 | -0.08 |
| *Gb_19468* | 0.05 | 0.11 | 363.23 | 254.56 | -0.08 |
| *Gb_40692* | 0.07 | 0.13 | 665.11 | 909.85 | -0.08 |
| *Gb_33207* | 0.05 | 0.08 | 459.27 | 396.18 | -0.08 |
| *Gb_34671* | 0.02 | 0.03 | 184.22 | 321.28 | -0.07 |

| | | | | | |
|---|---|---|---|---|---|
| *Gb_34669* | 0.02 | 0.03 | 184.22 | 321.29 | -0.07 |
| *Gb_05798* | 0.01 | 0.03 | 173.44 | 154.05 | -0.07 |
| *Gb_39329* | 0.00 | 0.01 | 65.42 | 53.30 | -0.07 |
| *Gb_15727* | 0.07 | 0.10 | 1402.73 | 1418.75 | -0.07 |
| *Gb_31200* | 27.64 | 33.41 | 0.00 | 0.01 | 0.07 |
| *Gb_29591* | 136.88 | 17.84 | 0.04 | 0.05 | 0.09 |
| *Gb_08147* | 8.64 | 1.86 | 0.00 | 0.01 | 0.09 |
| *Gb_16194* | 4.35 | 1.02 | 0.00 | 0.00 | 0.09 |
| *Gb_10867* | 21.13 | 7.34 | 0.01 | 0.03 | 0.09 |
| *Gb_09277* | 62.99 | 14.23 | 0.03 | 0.03 | 0.09 |
| *Gb_00826* | 75.08 | 29.25 | 0.04 | 0.13 | 0.09 |
| *Gb_27608* | 118.48 | 15.72 | 0.11 | 0.16 | 0.10 |
| *Gb_26510* | 35.37 | 7.87 | 0.03 | 0.07 | 0.10 |
| *Gb_11717* | 62.02 | 72.61 | 0.06 | 0.13 | 0.10 |
| *Gb_12430* | 3.77 | 1.28 | 0.00 | 0.01 | 0.10 |
| *Gb_19365* | 15.13 | 4.25 | 0.01 | 0.03 | 0.10 |
| *Gb_33039* | 47.21 | 10.11 | 0.05 | 0.10 | 0.10 |
| *Gb_31099* | 21.14 | 9.26 | 0.03 | 0.03 | 0.10 |
| *Gb_13573* | 15.96 | 8.16 | 0.02 | 0.03 | 0.10 |
| *Gb_36284* | 13.25 | 2.08 | 0.02 | 0.05 | 0.10 |
| *Gb_39256* | 5.26 | 0.78 | 0.01 | 0.02 | 0.11 |
| *Gb_15886* | 5.62 | 2.89 | 0.01 | 0.03 | 0.11 |
| *Gb_04443* | 14.93 | 3.19 | 0.03 | 0.09 | 0.11 |
| *Gb_16004* | 13.65 | 2.92 | 0.03 | 0.05 | 0.11 |
| *Gb_39245* | 18.04 | 3.48 | 0.04 | 0.13 | 0.11 |
| *Gb_41821* | 46.85 | 18.90 | 0.11 | 0.33 | 0.11 |
| *Gb_12165* | 6.65 | 1.38 | 0.02 | 0.05 | 0.12 |
| *Gb_01314* | 32.79 | 5.60 | 0.09 | 0.15 | 0.12 |
| *Gb_40207* | 7.09 | 3.50 | 0.02 | 0.04 | 0.12 |
| *Gb_00260* | 37.06 | 17.59 | 0.10 | 0.13 | 0.12 |
| *Gb_23442* | 17.74 | 13.30 | 0.06 | 0.10 | 0.12 |
| *Gb_35482* | 86.05 | 106.32 | 0.29 | 0.33 | 0.12 |
| *Gb_24791* | 28.18 | 4.53 | 0.10 | 0.19 | 0.12 |
| *Gb_19363* | 12.53 | 3.41 | 0.04 | 0.05 | 0.12 |
| *Gb_30992* | 15.58 | 3.74 | 0.05 | 0.09 | 0.12 |
| *Gb_26511* | 7.64 | 2.02 | 0.03 | 0.04 | 0.12 |
| *Gb_38310* | 7.43 | 5.02 | 0.03 | 0.04 | 0.12 |
| *Gb_24287* | 67.73 | 18.97 | 0.25 | 0.28 | 0.12 |
| *Gb_24902* | 8.93 | 6.42 | 0.03 | 0.05 | 0.12 |
| *Gb_17233* | 18.58 | 2.89 | 0.07 | 0.11 | 0.13 |
| *Gb_35476* | 55.60 | 72.85 | 0.23 | 0.16 | 0.13 |
| *Gb_11710* | 22.88 | 6.54 | 0.10 | 0.16 | 0.13 |
| *Gb_14342* | 32.13 | 3.83 | 0.16 | 0.19 | 0.13 |

| | | | | | |
|---|---|---|---|---|---|
| *Gb_22275* | 28.12 | 7.08 | 0.15 | 0.23 | 0.13 |
| *Gb_10165* | 18.29 | 6.26 | 0.11 | 0.09 | 0.13 |
| *Gb_40625* | 10.76 | 1.28 | 0.06 | 0.15 | 0.14 |
| *Gb_21651* | 9.37 | 2.13 | 0.06 | 0.08 | 0.14 |
| *Gb_24115* | 12.25 | 4.68 | 0.08 | 0.15 | 0.14 |
| *Gb_30534* | 34.14 | 4.88 | 0.22 | 0.26 | 0.14 |
| *Gb_12367* | 14.15 | 8.03 | 0.09 | 0.07 | 0.14 |
| *Gb_38764* | 19.66 | 3.17 | 0.13 | 0.19 | 0.14 |
| *Gb_37818* | 10.99 | 3.59 | 0.07 | 0.16 | 0.14 |
| *Gb_31928* | 6.56 | 1.15 | 0.04 | 0.06 | 0.14 |
| *Gb_22328* | 11.76 | 4.88 | 0.08 | 0.24 | 0.14 |
| *Gb_38308* | 7.32 | 4.11 | 0.05 | 0.11 | 0.14 |
| *Gb_27837* | 23.38 | 4.54 | 0.17 | 0.19 | 0.14 |
| *Gb_04442* | 17.98 | 3.81 | 0.14 | 0.22 | 0.14 |
| *Gb_16114* | 5.75 | 1.46 | 0.05 | 0.11 | 0.14 |
| *Gb_38662* | 28.14 | 4.12 | 0.23 | 0.38 | 0.14 |
| *Gb_23787* | 16.17 | 5.73 | 0.15 | 0.11 | 0.15 |
| *Gb_08907* | 26.60 | 2.76 | 0.24 | 0.34 | 0.15 |
| *Gb_20971* | 3.60 | 0.88 | 0.03 | 0.09 | 0.15 |
| *Gb_20908* | 27.91 | 7.43 | 0.26 | 0.34 | 0.15 |
| *Gb_34103* | 25.54 | 6.88 | 0.26 | 0.23 | 0.15 |
| *Gb_11719* | 12.04 | 8.26 | 0.13 | 0.12 | 0.15 |
| *Gb_24438* | 5.63 | 1.79 | 0.06 | 0.08 | 0.15 |
| *Gb_01779* | 3.16 | 0.59 | 0.03 | 0.06 | 0.15 |
| *Gb_04836* | 5.40 | 2.38 | 0.06 | 0.05 | 0.15 |
| *Gb_17798* | 14.71 | 3.41 | 0.17 | 0.27 | 0.16 |
| *Gb_01298* | 4.32 | 2.04 | 0.05 | 0.06 | 0.16 |
| *Gb_35711* | 14.21 | 4.50 | 0.17 | 0.19 | 0.16 |
| *Gb_13122* | 5.23 | 1.69 | 0.06 | 0.14 | 0.16 |
| *Gb_01435* | 4.14 | 1.82 | 0.05 | 0.11 | 0.16 |
| *Gb_14376* | 6.75 | 2.30 | 0.09 | 0.09 | 0.16 |
| *Gb_28308* | 11.83 | 6.96 | 0.17 | 0.20 | 0.16 |
| *Gb_33044* | 4.42 | 1.28 | 0.06 | 0.16 | 0.16 |
| *Gb_24464* | 14.74 | 3.99 | 0.23 | 0.33 | 0.17 |
| *Gb_24220* | 16.49 | 6.37 | 0.28 | 0.38 | 0.17 |
| *Gb_40750* | 3.34 | 0.70 | 0.06 | 0.06 | 0.17 |
| *Gb_30145* | 5.85 | 1.35 | 0.10 | 0.22 | 0.17 |
| *Gb_29836* | 9.84 | 3.89 | 0.18 | 0.25 | 0.17 |
| *Gb_20907* | 15.32 | 3.12 | 0.28 | 0.25 | 0.17 |
| *Gb_33551* | 16.57 | 3.19 | 0.30 | 0.29 | 0.17 |
| *Gb_00478* | 7.01 | 2.24 | 0.13 | 0.20 | 0.17 |
| *Gb_22074* | 6.29 | 1.37 | 0.12 | 0.18 | 0.17 |
| *Gb_28887* | 13.27 | 1.87 | 0.26 | 0.15 | 0.18 |

| | | | | | |
|---|---|---|---|---|---|
| *Gb_14457* | 23.62 | 3.10 | 0.49 | 0.27 | 0.18 |
| *Gb_29432* | 10.72 | 3.75 | 0.23 | 0.12 | 0.18 |
| *Gb_16734* | 5.61 | 1.68 | 0.12 | 0.17 | 0.18 |
| *Gb_12560* | 5.43 | 0.63 | 0.12 | 0.16 | 0.18 |
| *Gb_17118* | 8.85 | 1.71 | 0.20 | 0.17 | 0.18 |
| *Gb_33043* | 4.77 | 0.98 | 0.11 | 0.18 | 0.18 |
| *Gb_25645* | 11.72 | 3.53 | 0.28 | 0.27 | 0.19 |
| *Gb_16008* | 3.23 | 0.42 | 0.08 | 0.09 | 0.19 |
| *Gb_00316* | 6.63 | 2.83 | 0.18 | 0.25 | 0.19 |
| *Gb_09676* | 3.92 | 1.48 | 0.11 | 0.16 | 0.19 |
| *Gb_40268* | 9.97 | 2.83 | 0.28 | 0.26 | 0.19 |
| *Gb_38013* | 9.79 | 3.68 | 0.28 | 0.23 | 0.20 |
| *Gb_04740* | 9.00 | 1.42 | 0.26 | 0.20 | 0.20 |
| *Gb_33041* | 11.08 | 1.50 | 0.33 | 0.31 | 0.20 |
| *Gb_06085* | 11.05 | 1.36 | 0.33 | 0.33 | 0.20 |
| *Gb_17580* | 7.36 | 1.38 | 0.23 | 0.35 | 0.20 |
| *Gb_34690* | 7.76 | 4.41 | 0.25 | 0.20 | 0.20 |
| *Gb_38014* | 13.17 | 4.67 | 0.43 | 0.29 | 0.20 |
| *Gb_26697* | 5.57 | 1.72 | 0.18 | 0.28 | 0.20 |
| *Gb_06006* | 10.93 | 1.83 | 0.36 | 0.35 | 0.20 |
| *Gb_06280* | 3.33 | 0.70 | 0.11 | 0.17 | 0.20 |
| *Gb_08938* | 4.67 | 1.13 | 0.16 | 0.20 | 0.20 |
| *Gb_18580* | 3.54 | 0.56 | 0.12 | 0.15 | 0.21 |
| *Gb_23533* | 5.14 | 0.89 | 0.18 | 0.13 | 0.21 |
| *Gb_38060* | 5.62 | 1.94 | 0.20 | 0.24 | 0.21 |
| *Gb_02074* | 3.25 | 0.73 | 0.12 | 0.22 | 0.21 |
| *Gb_03444* | 4.47 | 1.21 | 0.17 | 0.21 | 0.21 |
| *Gb_39526* | 8.29 | 1.00 | 0.32 | 0.21 | 0.21 |
| *Gb_17753* | 3.62 | 1.11 | 0.14 | 0.12 | 0.21 |
| *Gb_26596* | 9.24 | 1.89 | 0.36 | 0.34 | 0.21 |
| *Gb_07887* | 6.23 | 1.91 | 0.25 | 0.22 | 0.22 |
| *Gb_39203* | 6.48 | 1.02 | 0.26 | 0.31 | 0.22 |
| *Gb_01720* | 4.87 | 0.43 | 0.21 | 0.27 | 0.22 |
| *Gb_26119* | 4.53 | 0.47 | 0.19 | 0.10 | 0.22 |
| *Gb_25388* | 7.81 | 1.40 | 0.34 | 0.17 | 0.22 |
| *Gb_01499* | 3.17 | 0.86 | 0.14 | 0.28 | 0.22 |
| *Gb_30001* | 4.13 | 0.61 | 0.19 | 0.15 | 0.22 |
| *Gb_22359* | 3.60 | 0.80 | 0.16 | 0.21 | 0.22 |
| *Gb_21015* | 4.20 | 0.80 | 0.20 | 0.28 | 0.23 |
| *Gb_11937* | 4.22 | 1.07 | 0.20 | 0.19 | 0.23 |
| *Gb_33755* | 5.07 | 1.98 | 0.26 | 0.28 | 0.23 |
| *Gb_16978* | 3.24 | 0.69 | 0.17 | 0.21 | 0.23 |
| *Gb_15873* | 6.65 | 1.47 | 0.36 | 0.23 | 0.24 |

| | | | | | |
|---|---|---|---|---|---|
| *Gb_36391* | 3.74 | 1.20 | 0.20 | 0.23 | 0.24 |
| *Gb_18105* | 9.05 | 1.20 | 0.52 | 0.27 | 0.24 |
| *Gb_11872* | 2.73 | 0.59 | 0.18 | 0.19 | 0.25 |
| *Gb_11136* | 7.38 | 1.14 | 0.48 | 0.30 | 0.25 |
| *Gb_19644* | 4.74 | 1.31 | 0.34 | 0.29 | 0.26 |
| *Gb_23886* | 4.01 | 1.06 | 0.30 | 0.29 | 0.27 |
| *Gb_26110* | 4.24 | 0.29 | 0.32 | 0.30 | 0.27 |
| *Gb_09024* | 3.72 | 0.58 | 0.29 | 0.27 | 0.27 |
| *Gb_39729* | 3.73 | 0.66 | 0.31 | 0.24 | 0.28 |
| *Gb_13711* | 5.82 | 1.04 | 0.49 | 0.26 | 0.28 |
| *Gb_22765* | 3.74 | 0.65 | 0.33 | 0.25 | 0.28 |
| *Gb_24439* | 3.08 | 0.66 | 0.28 | 0.25 | 0.29 |
| *Gb_20706* | 4.56 | 2.22 | 0.41 | 0.38 | 0.29 |
| *Gb_01837* | 3.11 | 0.81 | 0.29 | 0.33 | 0.29 |
| *Gb_16497* | 3.23 | 0.30 | 0.31 | 0.31 | 0.30 |
| *Gb_26313* | 3.08 | 0.63 | 0.33 | 0.41 | 0.31 |
| *Gb_30109* | 4.27 | 0.40 | 0.47 | 0.22 | 0.31 |
| *Gb_22956* | 3.40 | 0.64 | 0.39 | 0.35 | 0.32 |
| *Gb_29250* | 3.38 | 0.78 | 0.43 | 0.30 | 0.34 |
| *Gb_27802* | 2.77 | 0.55 | 0.36 | 0.35 | 0.34 |
| *Gb_14682* | 2.79 | 0.63 | 0.37 | 0.34 | 0.34 |
| *Gb_32086* | 2.48 | 0.28 | 0.37 | 0.17 | 0.36 |
| *Gb_15427* | 2.85 | 0.43 | 0.42 | 0.34 | 0.36 |
| *Gb_04723* | 2.83 | 0.56 | 0.48 | 0.20 | 0.39 |
| *Gb_12007* | 2.68 | 0.37 | 0.71 | 0.31 | 0.52 |
| *Gb_00825* | 45.42 | 24.75 | 0.00 | 0.00 | NULL |
| *Gb_04279* | 0.00 | 0.00 | 20.52 | 18.24 | NULL |
| *Gb_09785* | 4.97 | 1.22 | 0.00 | 0.00 | NULL |
| *Gb_10115* | 17.80 | 6.80 | 0.00 | 0.00 | NULL |
| *Gb_12438* | 5.71 | 1.74 | 0.00 | 0.00 | NULL |
| *Gb_14619* | 0.00 | 0.00 | 46.73 | 32.32 | NULL |
| *Gb_24705* | 2.45 | 0.49 | 0.00 | 0.00 | NULL |
| *Gb_25248* | 3.91 | 1.23 | 0.00 | 0.00 | NULL |
| *Gb_26840* | 3.87 | 0.94 | 0.00 | 0.00 | NULL |
| *Gb_26901* | 3.98 | 1.42 | 0.00 | 0.00 | NULL |
| *Gb_29390* | 6.51 | 1.50 | 0.00 | 0.00 | NULL |
| *Gb_32276* | 0.00 | 0.00 | 161.88 | 277.54 | NULL |
| *Gb_36507* | 4.62 | 1.30 | 0.00 | 0.00 | NULL |

**Table S8. Genes in the ginkgo sex-determination region from co-expression modules in the cone transcriptome.**

| No. | Gene ID | Module assigned |
|---|---|---|
| **1** | *Gb_15883* | turquoise |
| **2** | *Gb_15886* | turquoise |
| **3** | *Gb_28585* | turquoise |
| **4** | *Gb_28586* | turquoise |
| **5** | *Gb_28588* | turquoise |
| **6** | *Gb_28589* | turquoise |
| **7** | *Gb_13760* | turquoise |
| **8** | *Gb_30341* | turquoise |
| **9** | *Gb_30342* | turquoise |
| **10** | *Gb_30343* | turquoise |
| **11** | *Gb_30345* | turquoise |
| **12** | *Gb_15885* | red |
| **13** | *Gb_28587* | red |
| **14** | *Gb_15884* | brown |
| **15** | *Gb_30344* | cyan |
| **16** | *Gb_28584* | magenta |
| **17** | *Gb_13763* | tan |

**Table S9. SNPs in the sex-determination region of ginkgo. The numbers of SNPs of different categories are shown for S1, S2 and S3 are shown. CDS denotes coding sequence (exons).**

| | S1 | S2 | S3 | Total |
|---|---|---|---|---|
| All | 12425 | 2067 | 4672 | 19164 |
| Intergenic | 12358 | 1608 | 3567 | 17533 |
| CDS | 26 | 206 | 38 | 270 |
| Intronic | 41 | 253 | 1067 | 1361 |
